# Hierarchical clustering of gene-level association statistics reveals shared and differential genetic architecture among traits in the UK Biobank

**DOI:** 10.1101/565903

**Authors:** Melissa R. McGuirl, Samuel Pattillo Smith, Björn Sandstede, Sohini Ramachandran

## Abstract

Genome-wide association (GWA) studies have generally focused on a single phenotype of interest. Emerging biobanks that pair genotype data from thousands of individuals with phenotype data using medical records or surveys enable testing for genetic associations in each phenotype assayed. However, methods for characterizing shared genetic architecture among multiple traits are lagging behind. Here, we present a new method, Ward clustering to identify Internal Node branch length outliers using Gene Scores (WINGS), for characterizing shared and divergent genetic architecture among multiple phenotypes. The objective of WINGS (freely available at https://github.com/ramachandran-lab/PEGASUS-WINGS) is to identify groups of phenotypes, or “clusters”, that share a core set of genes enriched for mutations in cases. We show in simulations that WINGS can reliably detect phenotype clusters across a range of percent shared architecture and number of phenotypes included. We then use the gene-level association test PEGASUS with WINGS to characterize shared genetic architecture among 87 case-control and seven quantitative phenotypes in 349,468 unrelated European-ancestry individuals from the UK Biobank. We identify 10 significant phenotype clusters that contain two to eight phenotypes. One significant cluster of seven immunological phenotypes is driven by seven genes; these genes have each been associated with two or more of those same phenotypes in past publications. WINGS offers a precise and efficient new application of Ward hierarchical clustering to generate hypotheses regarding shared genetic architecture among phenotypes in the biobank era.

## 1 Introduction

Since the 2007 publication of the Wellcome Trust Case Control Consortium’s landmark genome-wide association (GWA) study of seven common diseases using 14,000 cases and 3,000 common controls, GWA studies have grown dramatically in scope. Much attention has been given to the increasing number of individuals sampled in GWA studies (198 studies to date have analyzed over 100,000 individuals, data accessed at https://www.ebi.ac.uk/gwas/docs/file-downloads on Jan. 5 2019), as well as to the challenges of interpreting and validating the statistically associated variants identified in large-scale studies (for recent examples, see [1, 2, 3, 4, 5, 6, 7]). However, as “mega-biobank” datasets (used here as by Huffman [5] to mean “a study with phenotype and genotype data on >100,000 individuals. . . rather than to the physical sample repository”) such as the UK Biobank [8] and BioVU at Vanderbilt University [9, 10] are interrogated by medical and population geneticists, there is comparatively less discussion surrounding approaches to analyze multiple phenotypes in a single genomic study.

In particular, a fundamental question mega-biobanks can answer is whether shared genetic architecture among multiple phenotypes is detectable using summaries of germline genetic variation. Pickrell et al. 2016 [11] explicitly tested for pleiotropy among 42 complex traits, focusing on identifying colocalized variants in GWA studies for pairs of traits (see also [12], which tests for colocalization between eQTLs and associated variants for the same trait). While phenome-wide association studies (PheWAS; [13, 14]) and multivariate GWA studies have tested for statistical association between variants and multiple phenotypes [15, 16, 17, 18, 19], these studies, including [11, 12] share the central challenge of single-phenotype GWA studies: they focus on single variants assumed to act independently, making results difficult to interpret biologically for any complex traits.

As large-scale GWA studies find statistically associated variants spread uniformly throughout the genome [2, 3, 20] and that effect sizes have reached diminishing returns [7], gene-level association tests [21, 22, 23] can offer insight into gene sets and pathways that are enriched for mutations in cases for a phenotype of interest. Gene-level association tests not only allow for different mutations to be associated with the phenotype of interest in different cases, but also generate biologically interpretable hypotheses regarding genetic interactions that the GWA framework ignores [24]. Despite this, gene-level association tests have rarely been brought to bear on multivariate GWA datasets. One approach was developed by Chang and Keinan (disPCA, [25]), who applied principal components analysis to a matrix of gene-level association scores to detect clusters of phenotypes in two dimensions. However, their dimensionality reduction of the gene score matrix ignored minor axes of variation across gene scores for ease of visualization and distances between phenotypes in PC space were difficult to interpret. Thus, identifying phenotypes significantly enriched for shared mutations in mega-biobanks remains an open challenge.

In this study, we present Ward clustering to identify Internal Node branch length outliers using Gene Scores (WINGS), a flexible method for (*i*) computationally detecting phenotype clusters based on genelevel association scores, and (*ii*) ranking phenotype clusters based on their levels of significance. Given gene-level association test statistics for multiple phenotypes as input, WINGS enables the detection of a “core set” of genes — that is, genes enriched for mutations in cases — across multiple phenotypes.

In order to identify genetic architectures shared across a set of phenotypes, we first use PEGASUS [26, 27] (see section 2.2 for more details) to assign a feature vector whose elements are gene-level association *p*-values scores, or “gene scores”, to each phenotype. Each such feature vector is an element of a high-dimensional vector space whose dimension is given by the number of genes included in the GWA study data. Given a list of *N* phenotypes, this approach therefore yields *N* feature vectors. The more significant genes two phenotypes share, the closer their features vectors will be. Choosing a norm on the vector space in which the feature vectors lie allows us to compute pairwise distances between any two feature vectors, resulting in an *N × N* matrix of pairwise distances – we note that different norms will result in different distance matrices, and we use this fact in this study to emphasize different parts of a feature vector when identifying clusters. Once a distance matrix has been computed, we can use clustering algorithms (in our case, Ward hierarchical clustering) to divide the set of phenotypes into disjoint groups that separate feature vectors based on their pairwise distances.

While hierarchical clustering algorithms have proven effective across a range of applications [28, 29, 30], the typical output of these clustering methods is a dendrogram illustrating the sequential formation of clusters starting with each cluster containing only a single data point and ending with a single cluster containing all of the data points. Consequently, it is unclear how to distinguish significant clusters from non-significant clusters and often this is done by choosing a single cutoff height in the dendrogram or predetermining the number of desired clusters [31, 32, 33]. WINGS, by contrast, implements a multi-step algorithm to systematically identify and rank significant clusters, described in detail in Section 2.

We evaluate the performance of WINGS in simulations under a variety of genetic architectures within phenotypes and shared among phenotypes. Lastly, we apply WINGS to identify significant phenotype clusters across 87 case-control phenotypes and 7 quantitative phenotypes assayed in 349,468 unrelated European-ancestry individuals in the UK Biobank.

## 2 Materials and Methods

### 2.1 UK Biobank data processing

1. Genotype and phenotype data from the UK Biobank release [8] were extracted (488,377 individuals, 784,256 variants) and filtered as follows:

a. Genotype data were extracted from the chrom*.cal files using the UK Biobank gconv tool
b. Phenotype data were taken from our application-specific csv file for application 22419
2. Only individuals who self-identified as white British were included in the study cohort (57,275 individuals removed)
3. All monomorphic variants were removed (19,189 variants removed)
4. Individuals identified by the UK Biobank to have high heterozygosity, excessive relatedness, or aneuploidy were removed (1,550 individuals removed)
5. Variants with a minor allele frequency less than 2.5% were not included (253,939 variants removed)
6. Only variants found to be Hardy-Weinberg Equilibrium (Fisher’s exact test *p* > 10^−6^) using plink 2.0 [34] were included (40,433 variants removed)
7. Variants with missingness greater than 1% were removed (60,523 variants removed)
8. Individuals with greater than 5% genotype missingness were removed (38 individuals removed)
9. Individuals who were third-degree relatives or closer were removed using the following process: One individual was removed at random from any pair of individuals with a kinship coefficient greater than 0.0442, calculated using KING (version 2.0; [35])

Following these QC steps, 349,468 individuals who self-identified as British and 410,172 variants remained for analysis. In order to control for population structure within the remaining cohort, principal component analysis (PCA) was performed using flashpca (version 2.0; [36]) on SNPs passing QC that were also in linkage equilibrium (SNPs with *r*^2^ > 0.1 removed, resulting in 104,834 SNPs for PCA).

We analyzed phenotypes in two stages. We selected an initial set of 26 case-control phenotypes based on phenotypes that had been previously analyzed in Shi et al. [2] and Pickrell et al. [11] that also had at least 100 cases in our cohort. Those phenotypes that did not have at least 100 cases in our cohort after QC were not included in the analysis (Table S1). A genome-wide association (GWA) study was performed for each of these 26 case-control phenotypes using plink2 [34] including age, sex, and the first five principal components as covariates to control for population structure.

We then expanded our analysis to include 61 additional case-control phenotypes and 7 quantitative phenotypes from the UK Biobank. These phenotypes were selected only if they had more than 1,000 cases in the analyzed cohort.

### 2.2 Overview of WINGS pipeline

For each of the phenotypes being jointly studied (either in simulations, as detailed in the next subsection, or in the UK Biobank), we used PEGASUS [26] to calculate gene-level association *p*-values for all autosomal genes in the human genome with at least one SNP within a +/−50kb window (17,651 genes). PEGASUS, developed by our group [26, 27], models correlation among genotypes in a region using linkage disequilibrium, the same model as VEGAS [21] and SKAT without weighting rare variants [22]. PEGASUS, by contrast, achieves up to machine precision in gene-level association statistic computations via numerical integration. In this study, we refer to the −log_10_ transformed PEGASUS gene-level association statistics as “gene-scores”.

Ward hierarchical clustering [37, 38] was then applied to the phenotypes using the PEGASUS gene scores (−log_10_ transformed PEGASUS *p*-values) as feature vectors. We then concatenate together each phenotype’s feature vector to generate a phenotype by gene matrix, the ultimate input for the WINGS software. In our analyses of the UK Biobank, a set of 7 continuous phenotypes were clustered separately due to their comparatively much larger sample sizes (Supplementary Figure S9 shows how the continuous and binary phenotypes cluster when treated as a single data set). Significant clusters were identified and ranked using the WINGS branch length thresholding algorithm (described in the Section 2.3).

### 2.3 WINGS, a new method for automatic phenotype cluster detection and ranking

WINGS is a thresholded hierarchical clustering algorithm that takes a matrix of gene-level association test results as its input and outputs identified phenotype clusters ranked by their significance.

First, WINGS applies Ward hierarchical clustering to the matrix of gene-level association test results, which we compute using PEGASUS. Specifically, consider a data set with *N* data points. Ward hierarchical clustering is an agglomerative clustering algorithm: initially there are *N* clusters, each containing exactly one data point, and clusters are merged recursively in a hierarchical manner until there is a single cluster containing all *N* data points [37, 38, 33].

Using an objective function approach, at each stage in an agglomerative clustering algorithm the pair of clusters that minimizes *the merging cost* are combined to form a single cluster. For Ward hierarchical clustering, the merging cost for combining clusters R and S of size *N_R_* and *N_S_* respectively, is defined as

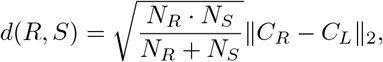

where *C_R_* and *C_L_* are the centroids of clusters R and L, respectively, and ‖ · ‖_2_ denotes the Euclidean norm. Note, this merging cost is equivalent to minimizing the increased sum of squared errors [37, 38, 33].

The choice to use Ward as the linkage criteria for WINGS was not arbitrary. Ward hierarchical clustering focuses on minimizing differences within the clusters, rather than maximizing pairwise distances between clusters. Previous work on comparing different agglomerative hierarchical clustering algorithms suggests that Ward clustering performs the best when clustering high dimensional, noisy data as long as cluster sizes are assumed to be approximately equal [39, 40]. We also note that we applied other linkage criteria to the data for comparison (see Supplement Section 5.3 and Supplement Figures S10-S15 for more details).

Hierarchical clustering results are often represented in a dendrogram, where each branch corresponds to a cluster, but it is not clear how to extract the clusters that are most significant [33, 31, 32]. Intuitively, significant clusters are those that form early on in the hierarchical clustering algorithm and do not merge with other clusters until there are very few clusters left. This corresponds to clusters that form near the bottom of the representative dendrogram tree and have long branch lengths.

To quantitatively define the notion of significantly long branch length we look at the consecutive differences between branch lengths and search for large gaps in the branch length distributions. That is, in the second step of WINGS we implement the following branch length thresholding algorithm to identify significant phenotype clusters within a dendrogram:

1. Sort all the branch lengths corresponding to small clusters (we define small clusters to be those with less than 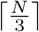 members, but the user can adjust this threshold);
2. Calculate the consecutive differences between branch lengths to get the branch length gaps;
3. Identify branch length gaps that are more than three scaled median absolute deviations away from the median and classify these as branch length gap outliers;
4. Set the branch length threshold to be the minimum branch length such that the branch length is greater than the median of all branch lengths and its branch length gap is a branch length gap outlier. If this threshold does not exist, we conclude that there are no significant clusters.

Finally, significant clusters are identified as the clusters whose corresponding dendrogram branch length is greater than or equal to the branch length threshold defined above.

Note that the branch length thresholding algorithm in WINGS is a multi-step process for identifying significant clusters in a dendrogram that does not require prior knowledge of the number of desirable clusters and is more flexible than the traditional fixed branch cut methods [32]. Previous work in [31] similarly introduces a dynamic method for identifying clusters from a dendrogram tree. In contrast to the iterative tree-cut algorithms presented in [31], however, WINGS relies solely on the dendrogram branch lengths and does not rely on making any tree cuts.

WINGS was implemented in MATLAB (R2017b) and applied to both simulated gene score matrices and empirical PEGASUS gene scores for phenotypes in the UK Biobank. These results are presented in the Section 3.

### 2.4 Simulations of phenotypes with shared genetic architecture

To test the sensitivity of WINGS when identifying both ground truth shared genetic architecture and varying levels of random noise in gene-level association *p*-values, we first applied WINGS to simulated gene score matrices. We also explored the differences between clusters identified by the raw PEGASUS *p*-values and clusters identified by the −log_10_ transformed PEGASUS *p*-values (“gene scores”). To accomplish these tasks, we generated both “significant genetic architectures”, where shared genes have a PEGASUS gene-level *p*-value < 0.001, and “non-significant architectures”, where clusters share genes with a PEGASUS gene-level *p*-value > 0.7.

Gene scores obtained as −log_10_ transformed PEGASUS gene-level *p*-values range from (0, ∞), where the highly significant genes have high transformed gene scores. We expect that clusters in this space are driven by shared significant genetic architecture — that is, traits that have a high percentage of shared significant genes — since these features contribute the most to the pairwise distances between phenotypes. If we instead study the raw (untransformed) PEGASUS gene-level *p*-values, we expect to see clusters of shared non-significant genetic architecture, referring to traits that have a high percentage of shared non-significant genes.

This distinction is illustrated in the synthetic example shown in Figure 1. As shown in Figure 1(A), groups of shared non-significant genetic architecture (shown in red) form clusters on the raw scale, whereas traits with shared significant genetic architecture (shown in orange) reside as a large, and therefore non-significant, group in the bottom left hand corner of the plot. In contrast, in Figure 1(B), groups of shared significant genetic architecture form clusters on the −log_10_ scale since this transformation maps the small region of significant *p*-values (gene-level *p* < 0.001) to the much larger region of (3, ∞).

**Figure 1:**
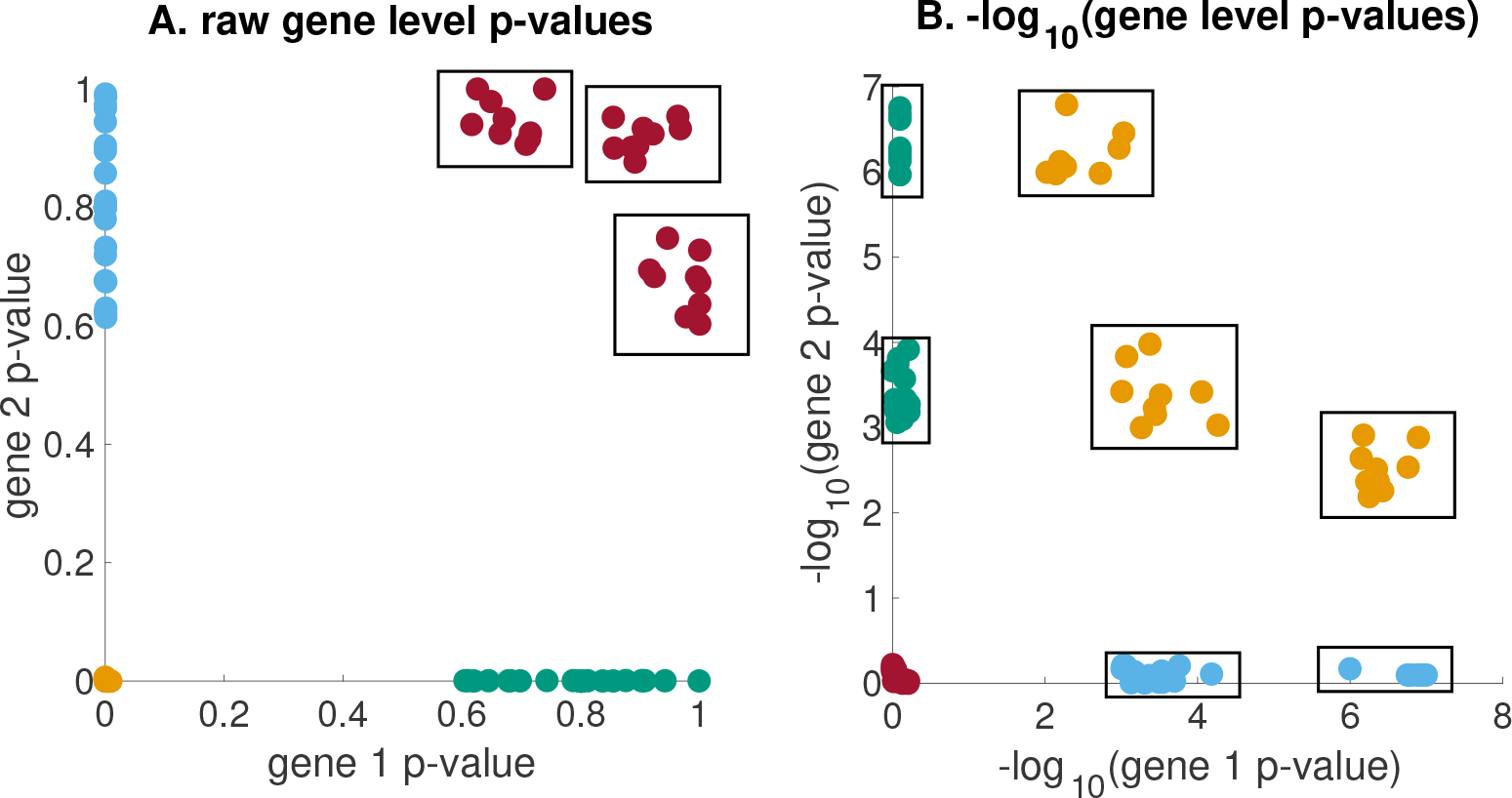
Synthetic clusters of traits with (A) shared non-significant genetic architecture and (B) shared significant genetic architecture from the raw and −log_10_ scales, respectively. Schematic example showing (A) simulated 2-dimensional gene-level *p*-values and (B) their corresponding −log_10_ transformed gene-level *p*-values. The boxed groups of points represent clusters of shared nonsignificant genetic architecture in (A) and clusters of shared significant genetic architecture in (B).

We now outline how we created simulated shared significant genetic architectures. Each simulated matrix was generated by randomly selecting PEGASUS *p*-values from the empirical distribution of PEGASUS *p*-values for Crohn’s disease (ICD10 code K50; 1,453 cases, 348,015 controls among the cohort passing our QC steps detailed earlier). PEGASUS *p*-values were then partitioned into significant (*p* < 0.001) and non-significant (*p* > 0.001) groups [41]. In the protocol described below, scores were taken randomly from the empirical gene scores in each of these groups. All simulated matrices maintain the same number of features (17,651 PEGASUS gene-level *p*-values, one for each autosomal gene) as our empirical analyses. For each phenotype in the matrix, 1% (175) of genes were assigned a significant value (*p* < 0.001).

We designed simulations that varied along two major parameters. We first set the number of phenotypes analyzed to either 25, 50, 75, or 100. Second, we set the percentage of the 175 significant genes that are shared between all cluster phenotypes to either 1% (2 genes), 10% (18 genes), 25% (44 genes), 50% (88 genes), or 75% (131 genes) as shared genetic architecture. For every pair of the parameters above we performed 1,000 simulations as detailed below.

In every simulation, the number and size of the clusters was determined using the following protocol:

1. Choose M from a uniform distribution between 3 and 15% of the total number of phenotypes; M will be the number of ground truth clusters simulated (e.g. for simulations with 100 phenotypes they all contain between 3 and 15 clusters)
2. For *j* = 1, 2, …, *M*

i. Generate ground truth cluster *j* of randomly selected phenotypes whose size is drawn at random from a uniform distribution between 2 and 8
ii. Select the corresponding percentage of significant genes to be shared for all phenotypes in the ground truth cluster
iii. Remove phenotypes in ground truth cluster *j* and corresponding shared significant genes from their respective pools (a phenotype may only be in one ground truth cluster, and a gene can only be shared and significant in one ground truth cluster)
iv. Assign non-shared significant genes and non-significant genes to each phenotype in the ground truth cluster
3. For all phenotypes not assigned to a ground truth cluster in Step 2, randomly draw 175 genes that remain in the pool to be significant and assign remaining genes as non-significant

Next, we focus on shared non-significant genetic architectures. We generated 1,000 additional simulations containing 75 phenotypes, using the same parameters for number of clusters and cluster size as described above, with the exception that we partitioned genes into those with PEGASUS gene-level *p*-values > 0.7 and those that have *p*-values < 0.7 and use the former to create clusters. Each of these simulations had the number of shared significant genes within a given cluster set to 75% (131 genes) of the 175 significant genes. We then analyzed each of the 1,000 simulations using the untransformed PEGASUS *p*-values and −log_10_ transformed data. We use the significant and non-significant genetic architecture simulations in tandem to better understand the driving factor of clusters identified by WINGS.

### 2.5 Gene Set Enrichment Analysis

Gene set enrichment analysis was performed for all significant empirical clusters. Genes that were significant (gene *p*-value < 0.001) for all cluster phenotypes were used as input into the EnrichR database [42]. The results from pathway analysis were used to annotate genes that WINGS identifies as associated with multiple traits.

### 2.6 Data Availability

Shared significant gene lists for each of the significant clusters in Figure 4, as well as scripts that were used to generate the simulated matrices and implement WINGS are available at https://github.com/ramachandran-lab/Pegasus-WINGS/.

## 3 Results

**Figure 2:**
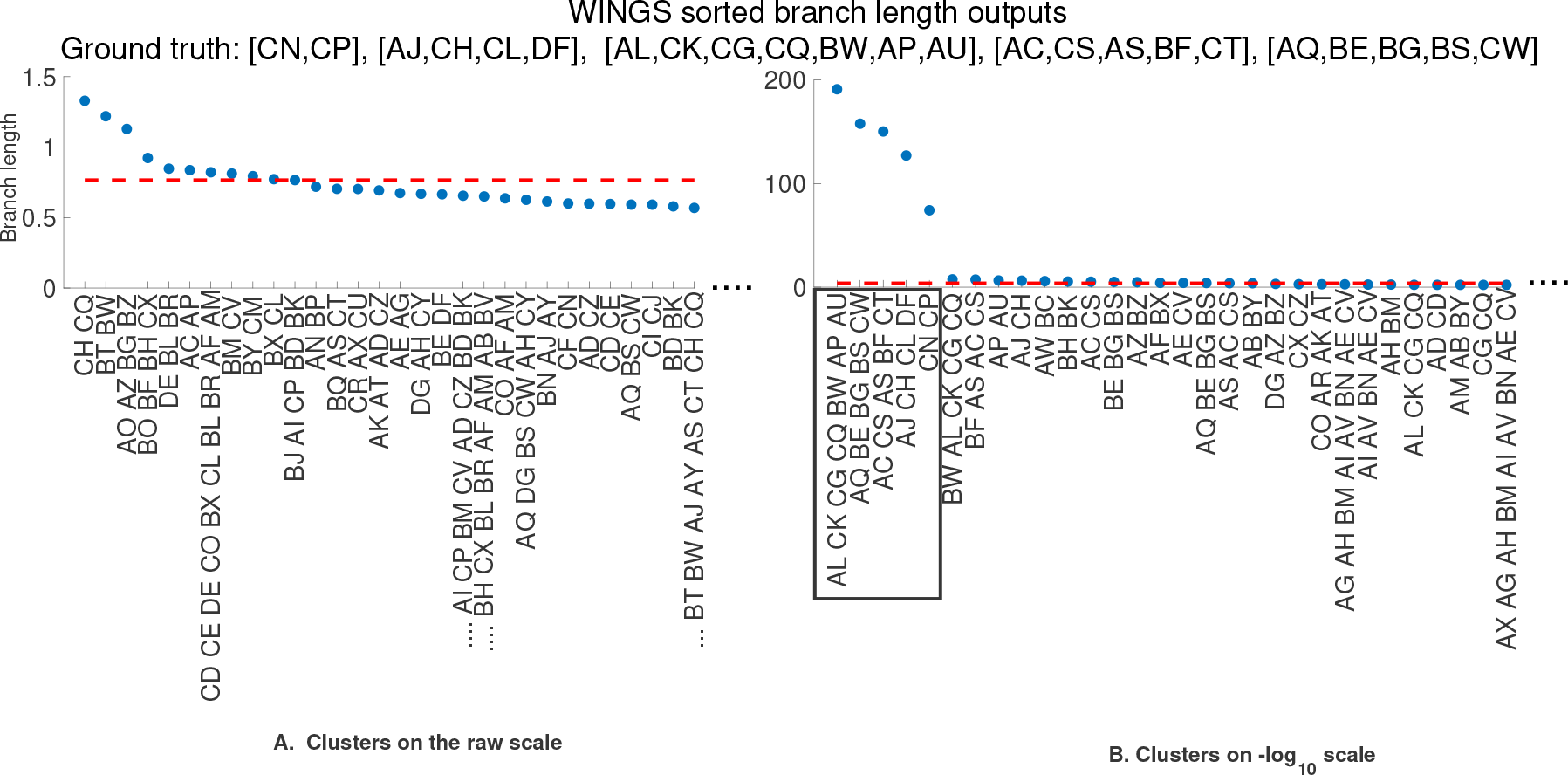
WINGS sorted branch lengths from a standard simulation identifies significant clusters on (A) raw and (B) −log_10_ scales. The sorted branch lengths corresponding to the dendrogram branches generated by WINGS when applied to the (A) raw PEGASUS gene-level *p*-values and (B) −log_10_ transformed PEGASUS gene scores from a simulation with 75 phenotypes, 75% (131) shared genes. For this simulation the ground truth clusters are [CN, CP], [AJ, CH, CL, DF], [AL, CK, CG, CQ, BW, AP, AU], [AC, CS, AS, BF, CT], and [AQ, BE, BG, BS, CW]. The dashed red horizontal line corresponds to the branch length threshold, where the identified significant clusters are those lying above the dashed line. The ground truth clusters are correctly identified as the significant clusters on the −log_10_ scale (boxed). These figures have been truncated on the right (removing some clusters that are not identified as significant) for better visualization purposes.

**Table 1:** Power (A) and precision (B) of WINGS across a range of phenotypes included as well as shared genetic architecture. “Shared genetic architecture” denotes the percentage of the 175 significant genes in each phenotype that are shared across all phenotypes in a cluster. Every entry in the table represents 1,000 simulations under the corresponding parameters. The power of WINGS for identifying ground truth clusters in simulations is defined as the percentage of ground truth clusters across these 1,000 simulations that were identified as significant by WINGS. The precision of WINGS is defined as follows: in a simulation with *x* ground truth clusters and a given number of phenotypes and proportion of shared genetic architecture, precision is the percentage of ground truth clusters that were identified as significant and within the *x* most significant clusters ranked by the branch thresholding step in WINGS.

| Phenotypes<br>in Simulation | Shared Genetic Architecture |  |  |  |  |  |  |  |  |  |
| --- | --- | --- | --- | --- | --- | --- | --- | --- | --- | --- |
|  | A: Power |  |  |  |  | B: Precision |  |  |  |  |
|  | 0.01 | 0.1 | 0.25 | 0.5 | 0.75 | 0.01 | 0.1 | 0.25 | 0.5 | 0.75 |
| 25 | 0.08 | 52.79 | 98.85 | 100 | 100 | 0.08 | 33.84 | 63.06 | 69.49 | 71.08 |
| 50 | 0.07 | 55.64 | 99.10 | 100 | 100 | 0.04 | 27.53 | 48.02 | 53.71 | 56.70 |
| 75 | 0.11 | 54.08 | 98.85 | 100 | 100 | 0.08 | 21.63 | 40.10 | 45.90 | 46.07 |
| 100 | 0.06 | 51.66 | 98.81 | 100 | 100 | 0.06 | 17.85 | 35.11 | 39.62 | 43.10 |

### 3.1 Performance on simulated data

In Table 1, we report power as the percentage of ground truth simulated clusters that WINGS correctly labels as significant across the 1,000 simulations, for a fixed number of phenotypes in analysis and percent shared significantly mutated genes (“shared genetic architecture”). We define shared genetic architecture for a cluster to be the percentage of genes that are significant (*p*-value < 0.001) across all member phenotypes of the cluster. We also measure the precision of WINGS in identifying simulated clusters. We define precision for a given simulation as the number of ground truth clusters that were correctly identified as significant and that further fell within the top *x* significant clusters in that simulation. For example, if a simulation has five ground truth clusters, the power of WINGS for that simulation would be the percentage of those five clusters that are identified as significant. The precision of WINGS is the percentage of those five ground truth clusters that have been both correctly identified and are within the five most significant clusters identified in that simulations. Table 1 reports the precision of WINGS on the standard simulations across varying parameter values for both the number of phenotypes analyzed and shared genetic architecture using PEGASUS *p*-values. We additionally generated simulations using the same protocol but substituting the PASCAL (sum) [23] and SKAT [43] gene-level association test results for PEGASUS gene-level association p-values to illustrate that WINGS can be used with any gene-level association metric. The results for the simulations using PASCAL and SKAT are shown in Table S2 and Table S3, respectively.

One sample output of WINGS applied to a standard simulation is presented in Figure 2. On the −log_10_ scale, the thresholded hierarchical clustering algorithm within WINGS identifies the ground truth clusters as the top five most significant clusters, whereas the clusters identified using raw gene-level *p*-values do not include the ground truth clusters. These results suggest that WINGS applied to −log_10_-transformed gene-level association statistics captures groups of phenotypes that have a high percentage of shared significant genes, but these ground truth clusters are not captured by the raw gene-level *p*-values.

**Table 2:** WINGS power and precision when applied to “non-significant architecture” simulations (see Methods, section 2.4); these simulations had 75 phenotypes and 75% (131 genes) shared genetic architecture.

|  | Power | Precision |
| --- | --- | --- |
| Raw PEGASUS $p$-values | 100 | 78.37 |
| $-\log_{10}$ PEGASUS $p$-values | 0.32 | 0.26 |

Using the protocol described in section 2.4, we applied WINGS to 1,000 non-significant architecture simulations to test its sensitivity to shared non-significant genetic architecture and analyzed the results. We find that WINGS is also robust to detecting shared levels of non-significant architecture using raw PEGASUS gene-level *p*-values (Table 2).

One sample output of WINGS applied to a non-significant architecture simulation with four ground truth clusters is presented in Figure 3. On the raw scale, the thresholded hierarchical clustering algorithm identifies the ground truth clusters as the top four most significant clusters, whereas the algorithm fails to identify the ground truth clusters when applied to the matrix of −log_10_ transformed PEGASUS gene-level *p*-values. These results suggest that clustering applied to raw PEGASUS gene-level *p*-values identifies clusters of phenotypes that have a high percentage of shared non-significant genes, while clustering using the −log_10_ transformed PEGASUS gene scores captures phenotype clusters that share a high percentage of significant genes.

### 3.2 Analysis of 87 case-control phenotypes

We first applied WINGS to the 26 case-control phenotypes analyzed in Pickrell et al. 2016 [11] and Shi et al. 2016 [2]. We provide the results of the analyzing these 26 phenotypes in 5. The focus of this paper is on the application of WINGS to 87 case-control phenotypes form the UK Biobank. We use the 26 phenotpypes from our initial analysis and add 61 case-control phenotypes that had at least 1,000 cases in our cohort from the UK Biobank (see Methods for QC details). The additional 61 phenotypes and their corresponding case numbers are provided in Table S1. We then applied WINGS to the resulting 87 phenotype by 17,651 gene matrix. In this expanded set of phenotypes, WINGS identifies 10 significant clusters, some of which contain smaller subclusters of phenotypes that are also significantly clustered. For instance, in Figure 4, the Immunological cluster 2 contains 7 phenotypes but many of the individual phenotype clades within it are additionally significant, including Type 1 diabetes mellitus (E10) and Seronegative rheumatoid arthritis (M06). For an exhaustive list of significant sub-clusters see Table 3. The ten significant clusters as well as their phenotypes are shown in the WINGS dendrogram in Figure 4 with the corresponding sorted branch length plots presented Figure S1. We find that the case number of a phenotype is not significantly correlated with that phenotype being in a significant cluster (Kendall’s *τ*, *p*-value < 0.2625). As expected, we found that whether a phenotype was in a significant cluster or not is significantly correlated with the number of significant gene scores (Kendall’s *τ*, *p*-value < 0.0002) when testing correlation between case number and number of significant PEGASUS gene scores after Bonferroni correction for 17,651 autosomal genes.

**Figure 3:**
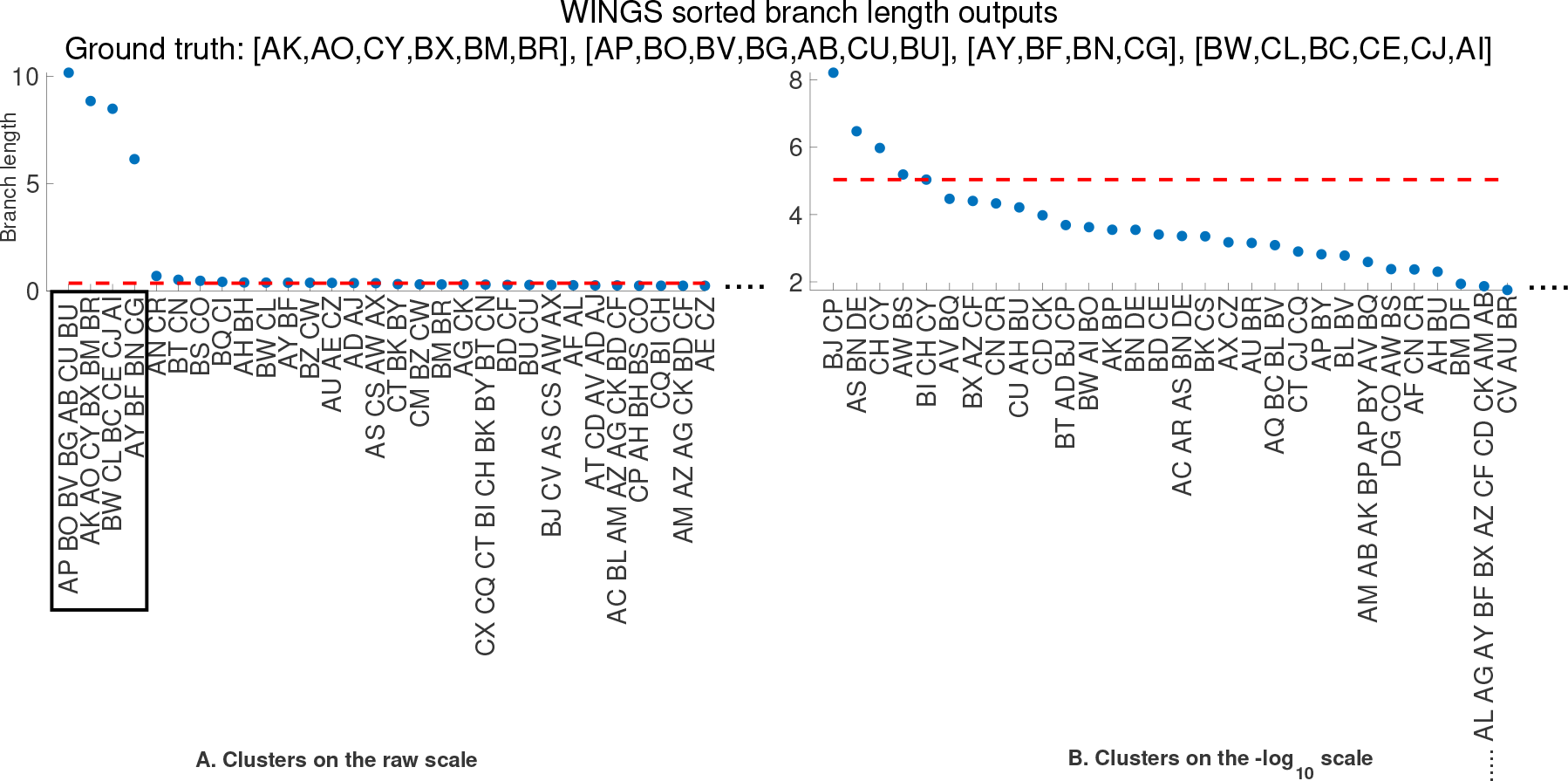
WINGS sorted branch lengths from a non-significant architecture simulation identifies significant clusters on (A) raw and (B) −log_10_ scales. Sorted branch lengths from the dendrogram output of WINGS when applied to (A) raw PEGASUS gene-level *p*-values and (B) −log_10_-transformed PEGASUS gene scores from a non-significant architecture simulation with 75 phenotypes and 75% shared non-significant genes. For this simulation the ground truth clusters are [AK, AO, CY, BX, BM, BR], [AP, BO, BV, BG, AB, CU, BU], [AY, BF, BN, CG], and [BW, CL, BC, CE, CJ, AI]. The dashed red horizontal line corresponds to the branch length threshold, above which the identified significant clusters lie. The ground truth clusters are correctly identified as the significant clusters on the raw scale (boxed). These figures have been truncated on the right (removing some clusters that are not identified as significant) for better visualization purposes.

In addition to only including genes and their +/− 50kb regions as features, we also computed PEGASUS scores for intergenic regions and observe that the topology of the tree is highly similar (dissimilarity index from [44] between Figure 4 and Figure S16 is Z = 0). We believe a more rigorous definition of intergenic regions may lead to a more informed tree.

### 3.3 Gene-Set Enrichment Analysis and Network Propagation

For each cluster, gene set enrichment analysis was performed using all genes that had a PEGASUS *p*-value < 0.001 [41] for every phenotype in a cluster. Significant genes shared by all phenotypes in the cluster of immunological phenotypes include several located in the MHC region: *BAT1*,*BAT3*,*BAT5* as well as *HLA-DOA* and *HLA-DRA* (See Supplementary Data on github for list of shared significant genes by cluster). Using the KEGG pathway database, Enrichr [45, 46] identified significant enrichment for genes that play a role in networks associated with Type I Diabetes mellitus, allograft rejection, and graft-versus-host disease.

**Figure 4:**
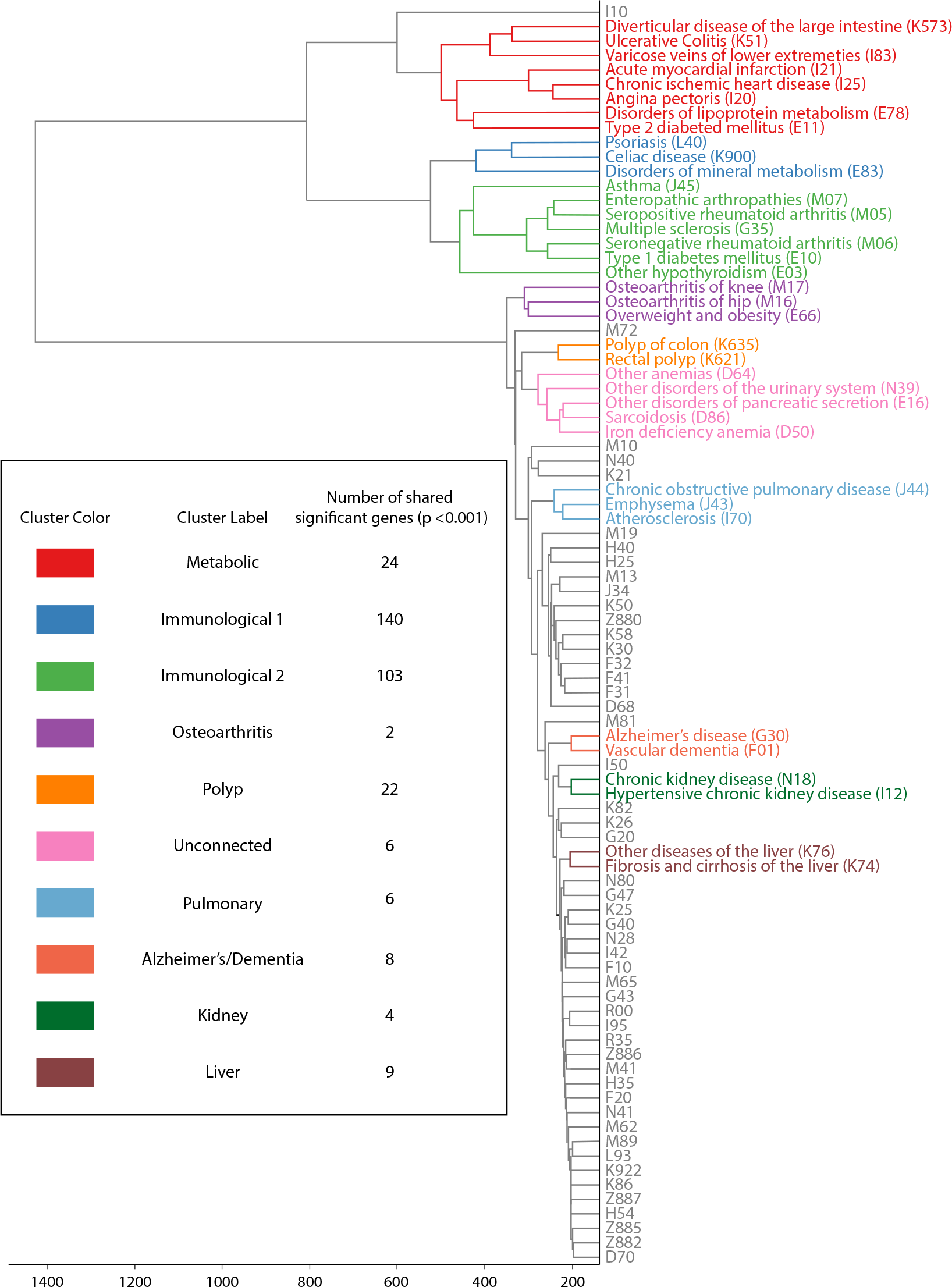
WINGS dendrogram from 87 case-control phenotypes in the UK Biobank reveals clusters of traits with shared significant genetic architecture. Dendrogram output from WINGS analysis of −log_10_ transformed PEGASUS scores of 87 case-control phenotypes in the UK Biobank. Listed are the ICD10 codes and common names of each phenotype that belongs to a significant cluster, grouped by cluster. Table insert: Each significant cluster’s color, assigned label, and number of shared significant genes (*p* < 0.001).

## 4 Discussion

Although biobank-scale datasets — in which multiple phenotypes are assayed and/or surveyed in tens of thousands to hundreds of thousands individuals — are becoming increasingly available to medical genomics researchers, approaches for leveraging these datasets to identify shared architecture among phenotypes are still in their infancy. Existing approaches for analyzing the shared genomic underpinnings of multiple phenotypes focus on colocalizing variant-level signals [14, 11], but these results overlook the role that genetic heterogeneity and interactions among genes may play in generating multiple complex traits and diseases.

Here, we present a new method, Ward clustering to identify Internal Node branch length outliers using Gene Scores (WINGS), for identifying phenotypes that share significant genetic architecture based on germline genetic data matched with binary or quantitative phenotypes for mega-biobanks. WINGS leverages Ward hierarchical clustering applied to gene scores for the phenotypes of interest, and further goes beyond past clustering applications to GWA studies of multiple phenotypes (e.g., disPCA; [25]) by providing a thresholding algorithm for identifying *significantly clustered* phenotypes. We note that the thresholding step in WINGS offers a useful visualization for interpreting results: while dendrograms depict the hierarchical architecture of clusters (Figures 4, S7-S8), the sorted branch lengths WINGS provides as output are intuitive to read, demonstrate a clear ranking of clusters, and identify significant clusters (Figures S1, S6).

Given concerns over whether GWA data contain signals of genetic architecture, we note that our simulations indicate that WINGS is sensitive to both *shared significant genes* (that is, genes enriched for trait-associated mutations) and *shared non-significant genes* (genes depleted for trait-associated mutations) (Figures 1-3; Tables 1,2). Figures 4, S6-S8 suggest that WINGS can offer insight into shared genetic architecture underlying comorbid phenotypes, or phenotypes that may often be misdiagnosed for one another such as vascular dementia and Alzheimer’s disease [47, 48]. Our results from applying WINGS to European-ancestry individuals sampled in the UK Biobank show that such relationships among phenotypes are not apparent from the taxonomy of ICD10 codes, where codes with the same letter prefix are considered related in their etiology.

Clustering of high-dimensional features will always be relative to the input data. In this case, an analysis of a subset of the phenotypes studied in the UK Biobank (Figures 4 and S8; Table S1) will alter results. Still, we underscore that our analysis of 26 phenotypes in the UK Biobank (chosen based on having been studied by both Pickrell et al. ([11]) and Shi et al. ([2]), as well as having over 100 cases in the UK Biobank) also recovers multiple significant clusters of phenotypes identified in our full set of 87 phenotypes: Alzheimer’s/Dementia, Metabolic and Immunological 2, Figure S8.

Next, we offer some caveats for future applications of WINGS and potential future directions for the development of methods to identify shared genetic architecture among multiple phenotypes in mega-biobanks. First, our goal here was to validate WINGS with simulations and to generate hypotheses regarding shared genetic architecture among complex traits in the UK Biobank. We do not seek to replicate our results from applying WINGS to data, an increasingly common challenge for mega-biobank analyses [5]. However, our validation with simulations and annotation of previously identified genes reinforces that we are reliably detecting true genetic architecture (see Supplementary File 1 for an extensive list of replication citations). Second, based on Figure S9, we do not suggest jointly studying traits with widely varying case numbers, in particular quantitative traits and binary phenotypes in mega-biobanks. One approach that could help overcome this challenge is the development of a gene score that incorporates both effect sizes and their standard errors into calculation [49], but this is outside the focus of our study.

Third, WINGS is sensitive to parameter choices: the clustering distance metric, the gene scores used as input, the upper-bound set for cluster size, and the branch length gap outlier criterion, which we will now discuss in more detail. We explored different clustering approaches beyond Ward hierarchical clustering using simulated data, and find that the choice of clustering method produces little change for results using raw gene-level *p*-values, but it does have a significant impact on clusters identified using −log_10_-transformed gene scores (Figures S10-S15). We focused on Ward hierarchical clustering here partly due to its performance on simulated phenotype clusters (Tables 1, 2), and due to its assumption that clusters are round; because clusters are hard to find in a high-dimensional space, this may be a conservative choice. We chose PEGASUS gene-level *p*-values as input to WINGS due to (*i*) our previous exploration of the power of PEGASUS ([26]); in particular PEGASUS is not biased by gene length, and computes more precise *p*-values than VEGAS [21] and SKAT [22]), and (*ii*) because the model of correlated SNP-level *p*-values underlying PEGASUS is the same as that of a number of gene-level association methods, Tables S2, S3. We set the upper bound on cluster size in our analyses to be *N*/3, where *N* is the number of phenotypes being analyzed, effectively discounting the potential for relatively large clusters, which we think is appropriate for analyses of mega-biobanks; future users may alter this threshold.

Future applications and extensions of WINGS may choose to explore a number of questions regarding shared genetic architecture among phenotypes. For example, [11] tested variants for true pleiotropy, while our current implementation of WINGS cannot differentiate between phenotypic relationships defined by clinical comorbidity versus causal dependence (see also [14]). We also assume that ICD10 codes are reliable indicators of disease status, which may not be the case [50, 51]. WINGS is sensitive to identifying shared mutated genes from −log_10_-transformed gene scores, and we interpret the genes underlying significant clusters in the output of WINGS as core genes underlying the clustered phenotypes [3]. Integrating results from WINGS with tissue-specific expression data would further test this hypothesis. WINGS could also be extended to test for differential genetic architecture among ancestries [52], a fundamental question to which mega-biobanks can offer unique insight in the coming years.

**Table 3:** Comparison of raw and −log_10_ significant phenotypes in the analysis of 87 case-control phenotypes. Clusters appearing on the same row have at least two common phenotypes in their intersection.

| Cluster Classification | Raw Scale | $-\log_{10}$ Scale |
| --- | --- | --- |
| Kidney Cluster | I12, N18 | I12, N18 |
| Liver Cluster | K74, K76 | K74, K76 |
| Polyp Cluster | K621, K635 | K621, K635 |
| Osteoarthritis Cluster | M13, M16, M17 | E66, M16, M17 |
| Metabolic Cluster | I20, I25<br>I20, I21, I25<br>E78, I10<br>E11, E78, I10<br>E11, E66, E78, I10, J45<br>E11, E66, E78, I10, I20, I21, I25, J45 | I20, I25<br>I20, I21, I25<br>E11, E78<br>E11, E78, I20, I21, I25<br>K51, K573<br>I83, K51, K573<br>E11, E78, I20, I21, I25, I83, K51, K573 |
| Immunology2 Cluster | M05, M06 | E10, M06<br>G35, M05, M07<br>E10, G35, M05, M06, M07<br>E10, G35, J45 M05, M06, M07<br>E03, E10, G35, J45 M05, M06, M07 |
| Unconnected Cluster | D50, D64 | D50, D86, E12<br>D50, D64, D86, E12, N39 |
| Immunology1 Cluster |  | K900, L40<br>E83, K900, L40 |
| Alzheimer’s/Dementia Cluster |  | F01, G30 |
| Pulmonary Cluster |  | I70, J43, J44 |
| No name | E03, M19 |  |
| No name | D70, G35 |  |
| No name | E83, Z882 |  |
| No name | L40, M07 |  |
| No name | J44, N39 |  |

## Acknowledgments

This research has been conducted using the UK Biobank Resource (Application #22419). Part of this research was conducted using computational resources and services at the Center for Computation and Visualization, Brown University. M.R.M. is supported by the National Science Foundation Graduate Research Fellowship Program under Grant No. 1644760. S.P.S. is a trainee in the Brown University Predoctoral Training Program in Biological Data Science, supported by NIH T32 GM128596. B.S. was partially supported by the National Science Foundation under grants DMS-1714429 and CCF-1740741. This work was also supported by US National Institutes of Health R01 GM118652 (to S.R.), and S.R. acknowledges additional support from National Science Foundation CAREER Award DBI-1452622. Any opinions, findings, and conclusions or recommendations expressed in this material are those of the authors and do not necessarily reflect the views of the National Science Foundation.

## 5 Supplemental Material

In this section we present the dendrogram outputs of WINGS applied to simulations presented in the main text and the branch length outputs applied to the −log_10_ transformed PEGASUS gene scores from 87 case-control phenotypes in the UK Biobank. Note, while the dendrograms contain information about the hierarchical architecture of the clusters, the sorted branch lengths presented in the paper are more intuitive to read, demonstrate a clear ranking of clusters, and identify the subset of highly significant clusters.

The branch length outputs outputs applied to the −log_10_ transformed PEGASUS gene scores from 87 case-control phenotypes in the UK Biobank. The corresponding dendrogram for this figure is presented in Figure 4 in the main text.

The dendrograms corresponding to clusters from the example simulation with 75% shared genes are presented in Figures S2-S3; and, the dendrograms corresponding to clusters from the example nonsignificant architecture simulation with 75% shared non-significant genes are presented in Figures S4-S5.

### Analysis of 26 case-control phenotypes

Here we present results from applying WINGS to 26 binary chronic illness phenotypes in the UK Biobank. Figure S6 displays the branch length outputs of WINGS (see Methods, section 2) applied to the raw and −log_10_-transformed PEGASUS gene scores computed using cases and controls from the UK Biobank for 26 binary chronic illness phenotypes that were also studied by Shi et al. [2] and Pickrell et al. [11].

On the raw scale, Figure S6(A) reveals that the significant clusters identified are [J45, E11, 125, E78], [M07, L40], and [M05, M06]. The significant −log_10_ clusters identified by WINGS in Figure S6(B) can be annotated as metabolic [E11, I25, E78], immunological [K900, J45, K51, L40, M06, G35, M05, M07], and Alzheimer’s/dementia [G30, F01] (see Table S1 for common disease names, as well as the shared significant genes in a cluster). On both scales, the clusters identified from WINGS applied to these 26 phenotypes in the UK Biobank are similar to the clusters identified from WINGS applied to 87 case-control phenotypes in the UK Biobank (see Table 3 and Figure 4 in the main text).

The dendrogram corresponding to clusters from the 26 phenotypes from the UK Biobank is presented in Figure S7. Figure S8 displays the dendrogram output of WINGS applied to the −log_10_-transformed PEGASUS gene scores for these 26 binary chronic illness phenotypes in the UK Biobank. The dendrogram displays the hierarchical nature of the immunological cluster (orange branches in Figure S8), and it demonstrates the proximity of the [G30, F01] cluster to other phenotypes.

### WINGS applied to 87 continuous and 7 binary phenotypes in the UK Biobank

Figure S9 displays results from simultaneously applying WINGS to 7 continuous and and 87 binary phenotypes. The binary traits and continuous traits cluster separately with the exception of nucleated red blood cells (NRB). We note that the NRB phenotype is only partially continuous in that there is a continuous spectrum of nucleated red blood cells for unhealthy individuals, but all healthy individuals will have a zero value. Thus, it is not surprising that NRB trait does not belong to a significant cluster.

Ignoring the NRB trait, the cluster of continuous phenotypes (represented in yellow on the far left of the dendrogram in Figure S9) remains completely disjoint from the discrete traits until there is only a single cluster containing all traits. We observe that the [BMI, WHR] cluster has 6,781 shared significant genes (*p* < 0.001); the [PLC, MCV, MCV] cluster has 3,553 shared significant genes; and, the full continuous cluster with traits [BMI, WHR, PLC, MCV, MCV, Height] has 1,685 shared significant genes. This is unsurprising as complex continuous phenotypes have been shown to be highly polygenic [3, 53, 54].

### Robustness to clustering criterion

In this paper, we present WINGS, a thresholded clustering algorithm based on Ward Hierarchical Clustering. While the Ward linkage criterion works well to cluster phenotypes, other linkage criterion may be used. To test the robustness of WINGS with respect to the choice of linkage criterion, we applied our method using single linkage, average linkage, and complete linkage clustering to the 26 phenotypes we analyzed from the UK Biobank in Section 5 (see [33] for more information on single linkage, average linkage, and complete linkage clustering). Here, we used the same branch thresholding algorithm described in Section 2.3 with each linkage criterion to identify significant clusters. For reference, the Ward-based WINGS results are presented in Figures S6-S8.

We observe that the significant clusters remain robust with respect to the linkage criterion when using raw PEGASUS gene-level *p*-values. When applied to −log_10_-transformed PEGASUS gene scores, however, the clusters appear to be more sensitive to the choice of linkage criterion. Future studies will be dedicated to fully understanding the differences between the clusters identified by WINGS, single linkage clustering, average linkage clustering, and complete linkage clustering on the −log_10_ scale.

The dendrograms and sorted branch length plots for these results are demonstrated in Figures S10-S15.

### WINGS clustering of 87 phenotypes using all genomic regions

In order to test if additional information about shared genetic architecture across phenotypes exists in intergenic region we performed an additional analysis. Using the bounds of the 17,651 genes (accounting for overlap) in our initial analysis to define 2,961 intergenic regions that were not included in the initial analysis. For each of these regions we performed a PEGASUS gene-level association test to generate a *p*-value for the region. We then pooled the *p*-values from our initial analysis with those of the intergenic regions to create a matrix of 87 phenotypes with 20,116 features (regional statistics). The resulting tree topology is shown in S16.

## Supplemental Tables and Figures

**Table S1:** Phenotypes analyzed in this study sorted by International Classication of Disease (ICD10) codes. * denotes that the phenotype was included in the initial analysis of 26 case-control traits that were also studied by Pickrell et al. [11] and Shi et al. [2]

| Disease | ICD10 Code | Number of Cases |
| --- | --- | --- |
| Iron deficiency anemia | D50 | 6284 |
| Other anemias | D64 | 9522 |
| Other coagulation defects | D68 | 809 |
| Neutropenia | D70 | 2636 |
| *Sarcoidosis | D86 | 449 |
| Other hypothyroidism | E03 | 11691 |
| Type 1 diabetes mellitus | E10 | 2373 |
| *Type 2 diabetes mellitus | E11 | 15080 |
| Other disorders of pancreatic internal secretion | E16 | 764 |
| Overweight and obesity | E66 | 8950 |
| *Disorders of lipoprotein metabolism and other lipidemias | E78 | 29778 |
| Disorders of mineral metabolism | E83 | 1758 |
| *Vascular dementia | F01 | 156 |
| Alcohol related disorders | F10 | 4313 |
| *Schizophrenia | F20 | 425 |
| *Bipolar disorder | F31 | 791 |
| *Major depressive disorder | F32 | 9714 |
| Other anxiety disorders | F41 | 4881 |
| *Parkinson's disease | G20 | 972 |
| *Alzheimer's disease | G30 | 331 |
| *Multiple sclerosis | G35 | 1124 |
| Epilepsy and recurrent seizures | G40 | 3071 |
| *Migraine | G43 | 2263 |
| Sleep disorders | G47 | 4410 |
| Age-related cataract | H25 | 6814 |
| Other retinal disorders | H35 | 2872 |
| Glaucoma | H40 | 3729 |
| Blindness and low visions | H54 | 728 |
| Hypertension | I10 | 64135 |
| Hypertensive chronic kidney disease | I12 | 1274 |
| Angina pectoris | I20 | 15063 |
| Acute myocardial infarction | I21 | 6655 |
| *Chronic ischemic heart disease | I25 | 20958 |
| Cardiomyopathy | I42 | 1035 |
| Heart failure | I50 | 4423 |
| Atherosclerosis | I70 | 1025 |
| Varicose veins of lower extremities | I83 | 8988 |
| Hypotension | I95 | 4072 |
| Other and unspecified disorders of nose and nasal sinuses | J34 | 5393 |
| Emphysema | J43 | 1388 |
| Other chronic obstructive pulmonary disease | J44 | 6833 |
| *Asthma | J45 | 21758 |
| Gastro-esophageal reflux disease | K21 | 19132 |
| Gastric ulcer | K25 | 3467 |
| Duodenal ulcer | K26 | 2517 |
| Functional dyspepsia | K30 | 9696 |
| *Crohn's disease | K50 | 1436 |
| *Ulcerative colitis | K51 | 2661 |
| Diverticular disease of large intestine without perforation or abscess | K573 | 19462 |
| Irritable bowel syndrome | K58 | 4563 |
| Rectal polyp | K621 | 5210 |
| Polyp of colon | K635 | 9306 |
| Fibrosis and cirrhosis of liver | K74 | 676 |
| Other diseases of liver | K76 | 2791 |
| Other diseases of gallbladder | K82 | 1482 |
| Other diseases of pancreas | K86 | 896 |
| *Celiac disease | K900 | 1522 |
| Gastrointestinal hemorrhage | K922 | 4387 |
| *Psoriasis | L40 | 1836 |
| *Lupus erythematosus | L93 | 105 |
| *Rheumatoid arthritis with rheumatoid factor | M05 | 465 |
| *Other rheumatoid arthritis | M06 | 3581 |
| *Enteropathic arthropathies | M07 | 591 |
| Gout | M10 | 2661 |
| Other arthritis | M13 | 9500 |
| Osteoarthritis of hip | M16 | 9876 |
| Osteoarthritis of knee | M17 | 16612 |
| Other and unspecified osteoarthritis | M19 | 13548 |
| Scoliosis | M41 | 838 |
| Other disorders of muscle | M62 | 746 |
| Synovitis and tenosynovitis | M65 | 4311 |
| Fibroblastic disorders | M72 | 3267 |
| Osteoporosis | M81 | 4884 |
| Other disorders of bone | M89 | 1261 |
| Chronic kidney disease | N18 | 3714 |
| Other disorders of kidney and ureter | N28 | 1996 |
| Other disorders of urinary system | N39 | 15870 |
| Benign prostatic hyperplasia | N40 | 9471 |
| Inflammatory diseases of prostate | N41 | 1334 |
| Endometriosis | N80 | 3235 |
| Abnormalities of heart beat | R00 | 7018 |
| Polyuria | R35 | 3191 |
| *Allergy status to penicillin | Z880 | 13436 |
| *Allergy status to sulfonamides status | Z882 | 712 |
| *Allergy status to narcotic agent status | Z885 | 983 |
| *Allergy status to analgesic agent status | Z886 | 3586 |
| *Allergy status to serum and vaccine status | Z887 | 157 |

**Table S2:** WINGS performance on simulated data generated using the empirical distribution of PASCAL [23] sum gene scores for Crohn’s disease (17,582 genes). Power (A) and precision (B) of WINGS across a range of phenotypes included as well as shared genetic architecture. “Shared genetic architecture” denotes the percentage of the 175 significant genes in each phenotype that are shared across all phenotypes in a cluster. Every entry in the table represent 1,000 simulations under the corresponding parameters. Power and precision are defined explicitly in Table 1.

| Phenotypes<br>in Simulation | Shared Genetic Architecture |  |  |  |  |  |  |  |  |  |
| --- | --- | --- | --- | --- | --- | --- | --- | --- | --- | --- |
|  | **A: Power** |  |  |  |  | **B: Precision** |  |  |  |  |
|  | 0.01 | 0.1 | 0.25 | 0.5 | 0.75 | 0.01 | 0.1 | 0.25 | 0.5 | 0.75 |
| 25 | 0.08 | 77.08 | 99.92 | 100 | 100 | 0.04 | 51.31 | 71.22 | 75.30 | 77.56 |
| 50 | 0.11 | 78.08 | 99.98 | 100 | 100 | 0.07 | 38.57 | 57.14 | 62.14 | 65.36 |
| 75 | 0.12 | 76.74 | 99.97 | 100 | 100 | 0.07 | 31.20 | 51.36 | 53.97 | 56.73 |
| 100 | 0.04 | 74.99 | 99.98 | 100 | 100 | 0.04 | 28.17 | 46.33 | 50.66 | 54.12 |

**Table S3:** WINGS performance on simulated data generated using the empirical distribution of SKAT [43] sum gene scores for Crohn’s disease (11,518 genes). Power (A) and precision (B) of WINGS across a range of phenotypes included as well as shared genetic architecture. “Shared genetic architecture” denotes the percentage of the 175 significant genes in each phenotype that are shared across all phenotypes in a cluster. Every entry in the table represent 1,000 simulations under the corresponding parameters. Power and precision are defined explicitly in Table 1.

| Phenotypes<br>in Simulation | Shared Genetic Architecture |  |  |  |  |  |  |  |  |  |
| --- | --- | --- | --- | --- | --- | --- | --- | --- | --- | --- |
|  | **A: Power** |  |  |  |  | **B: Precision** |  |  |  |  |
|  | 0.01 | 0.1 | 0.25 | 0.5 | 0.75 | 0.01 | 0.1 | 0.25 | 0.5 | 0.75 |
| 25 | 0.01 | 12.94 | 91.43 | 99.96 | 100 | 0.04 | 8.63 | 61.65 | 74.44 | 77.59 |
| 50 | 0.02 | 9.11 | 90.79 | 99.95 | 100 | 0.02 | 5.37 | 48.47 | 63.24 | 66.73 |
| 75 | 0.01 | 7.19 | 90.14 | 99.97 | 100 | 0.01 | 3.47 | 42.06 | 57.78 | 61.07 |
| 100 | 0.01 | 6.74 | 89.41 | 99.95 | 99.99 | 0.01 | 3.00 | 35.35 | 53.89 | 58.69 |

**Figure S1:**
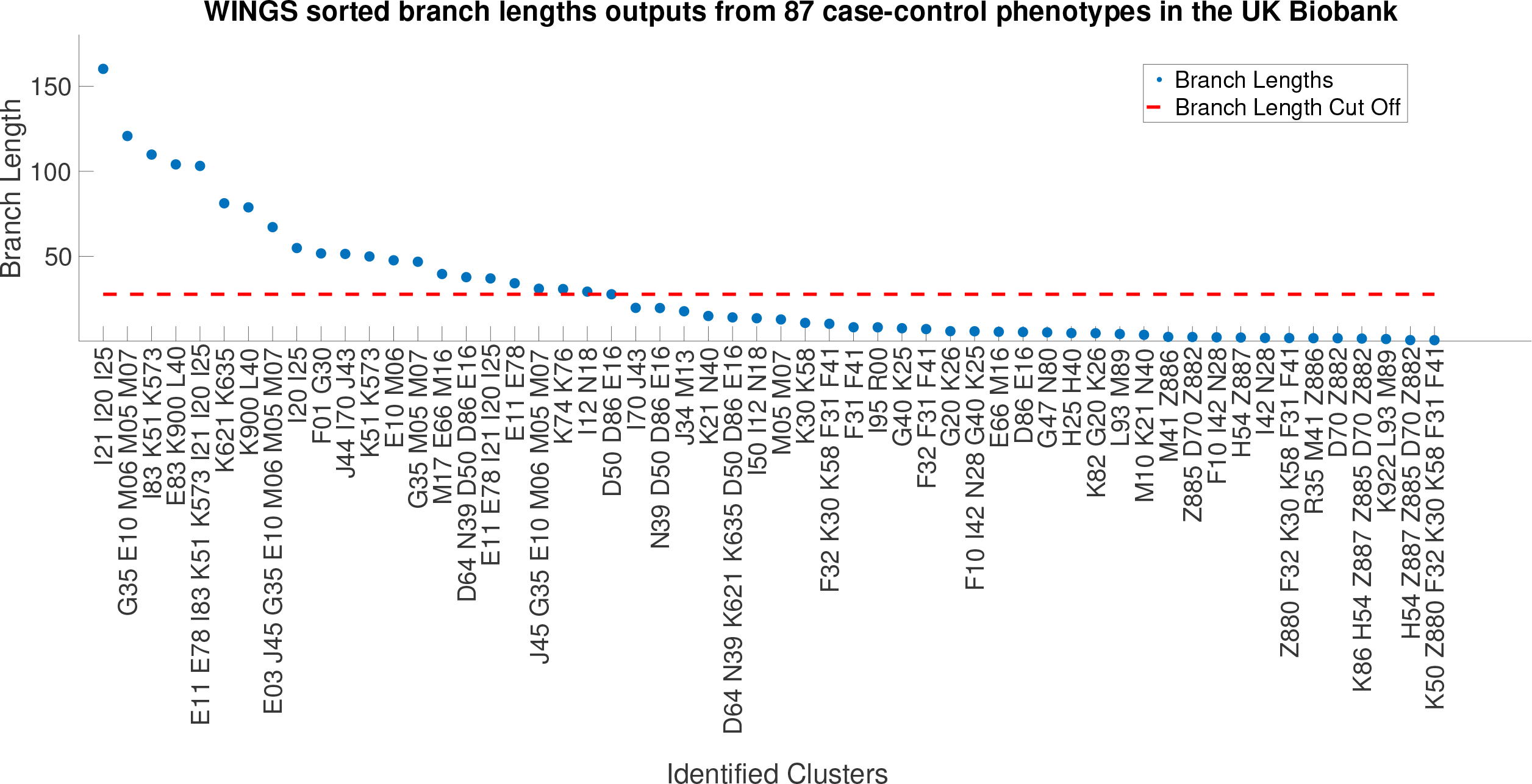
WINGS sorted branch lengths from 87 case-control phenotypes in the UK Biobank reveals clusters of traits with shared significant genetic architecture. The sorted branch lengths corresponding to the dendrogram branches generated by WINGS when applied to the −log_10_ transformed PEGASUS gene scores from 87 case-control phenotypes in the UK Biobank. The dashed red horizontal line corresponds to the branch length threshold, where the identified significant clusters are those lying above the dashed line. The corresponding dendrogram is presented in Figure 4.

**Figure S2:**
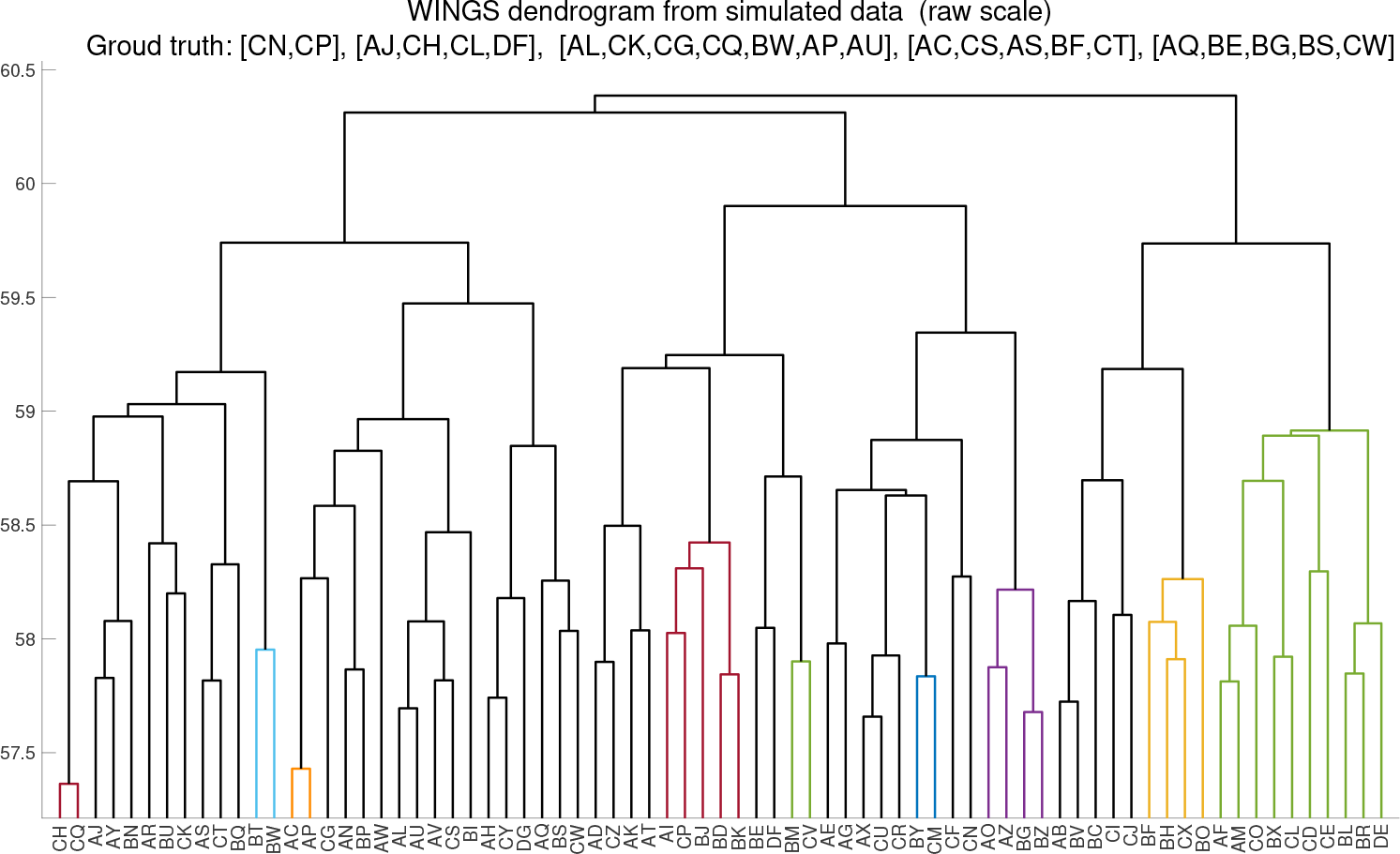
WINGS dendrogram from a standard simulation on the raw scale. The dendrogram output of Ward hierarchical clustering applied to the raw PEGASUS scores of a simulation with 75 traits, 75% shared genes. The branches are color coded by the largest significant clusters identified by the branch thresholding algorithm. The corresponding sorted branch lengths are presented in Figure 2 in the paper.

**Figure S3:**
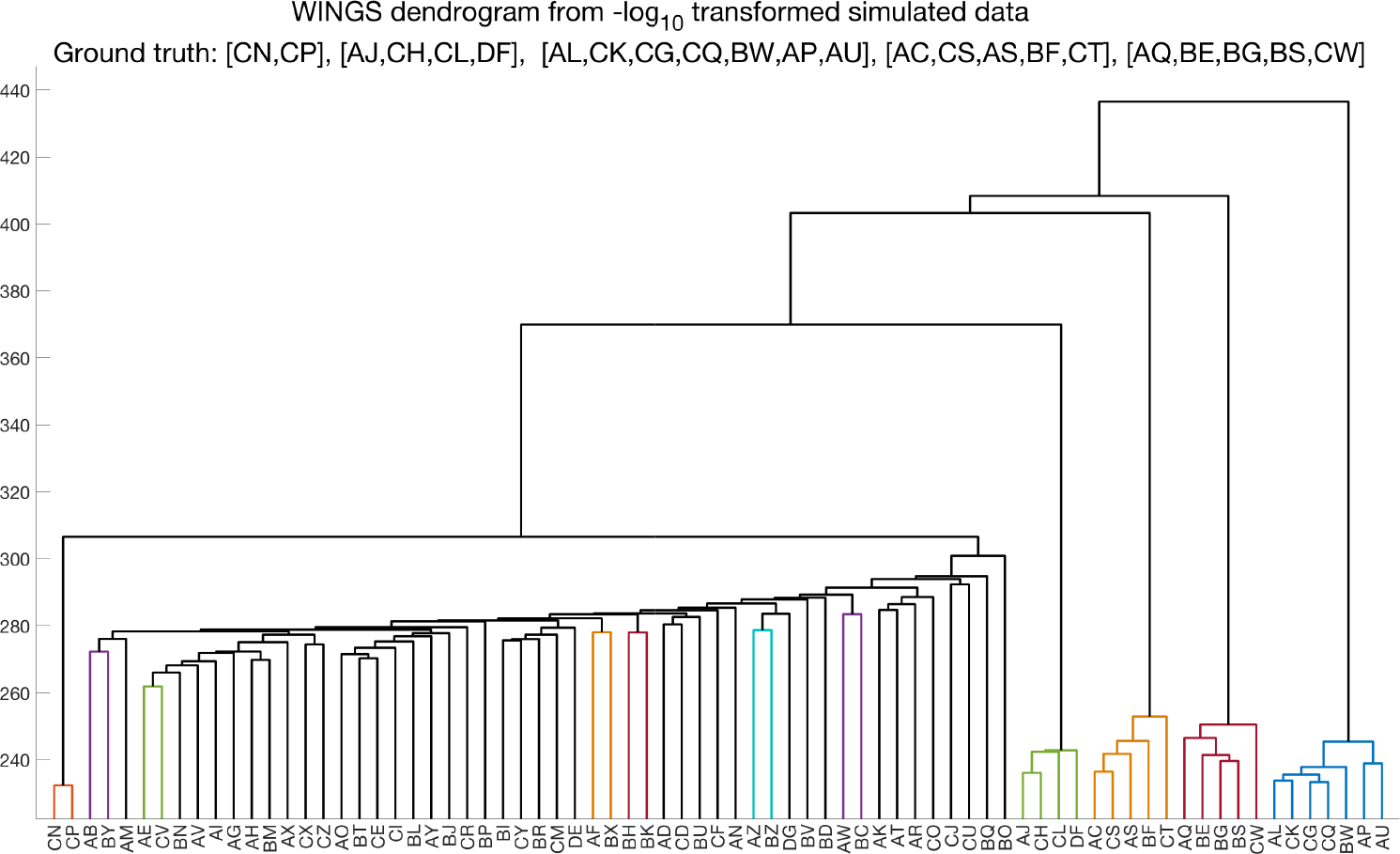
WINGS dendrogram from a standard simulation on the −log_10_ scale. The dendrogram output of Ward hierarchical clustering applied to the −log_10_ transformed PEGASUS scores of a simulation with 75 traits and 75% shared genes. The branches are color coded by the largest significant clusters identified by the branch thresholding algorithm. The corresponding sorted branch lengths are presented in Figure 2 in the paper.

**Figure S4:**
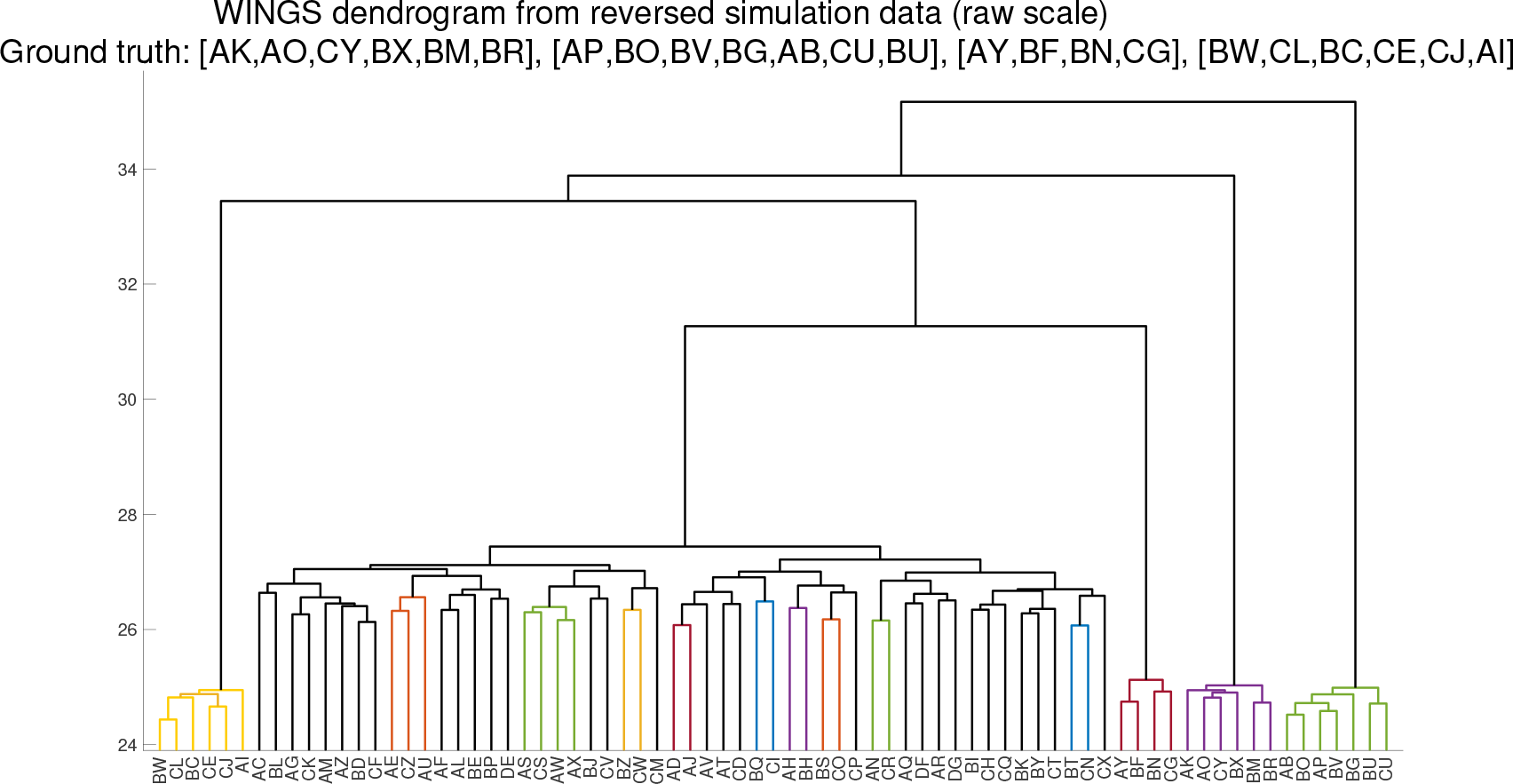
WINGS dendrogram from a non-significant architecture simulation on the raw scale. The dendrogram output of Ward hierarchical clustering applied to the raw PEGASUS scores of a non-significant architecture simulation with 75 traits and 75% shared genes. The branches are color coded by the largest significant clusters identified by the branch thresholding algorithm. The corresponding sorted branch lengths are presented in Figure 3 in the paper.

**Figure S5:**
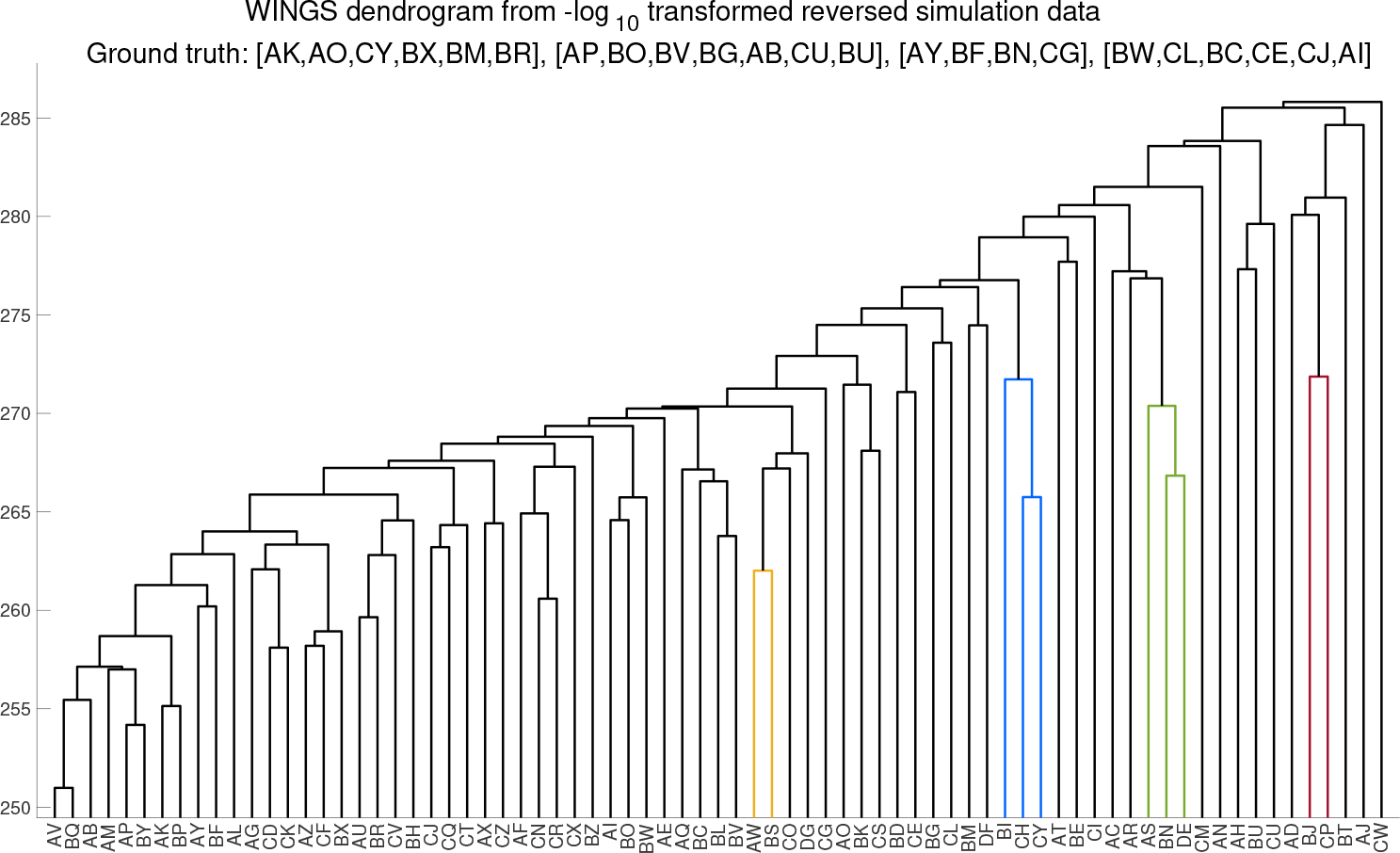
WINGS dendrogram from a non-significant architecture simulation on the −log_10_ scale. The dendrogram output of Ward hierarchical clustering applied to the −log_10_ transformed PEGASUS scores of a non-significant architecture simulation with 75 traits and 75% shared genes. The branches are color coded by the largest significant clusters identified by the branch thresholding algorithm. The corresponding sorted branch lengths are presented in Figure 3 in the paper.

**Figure S6:**
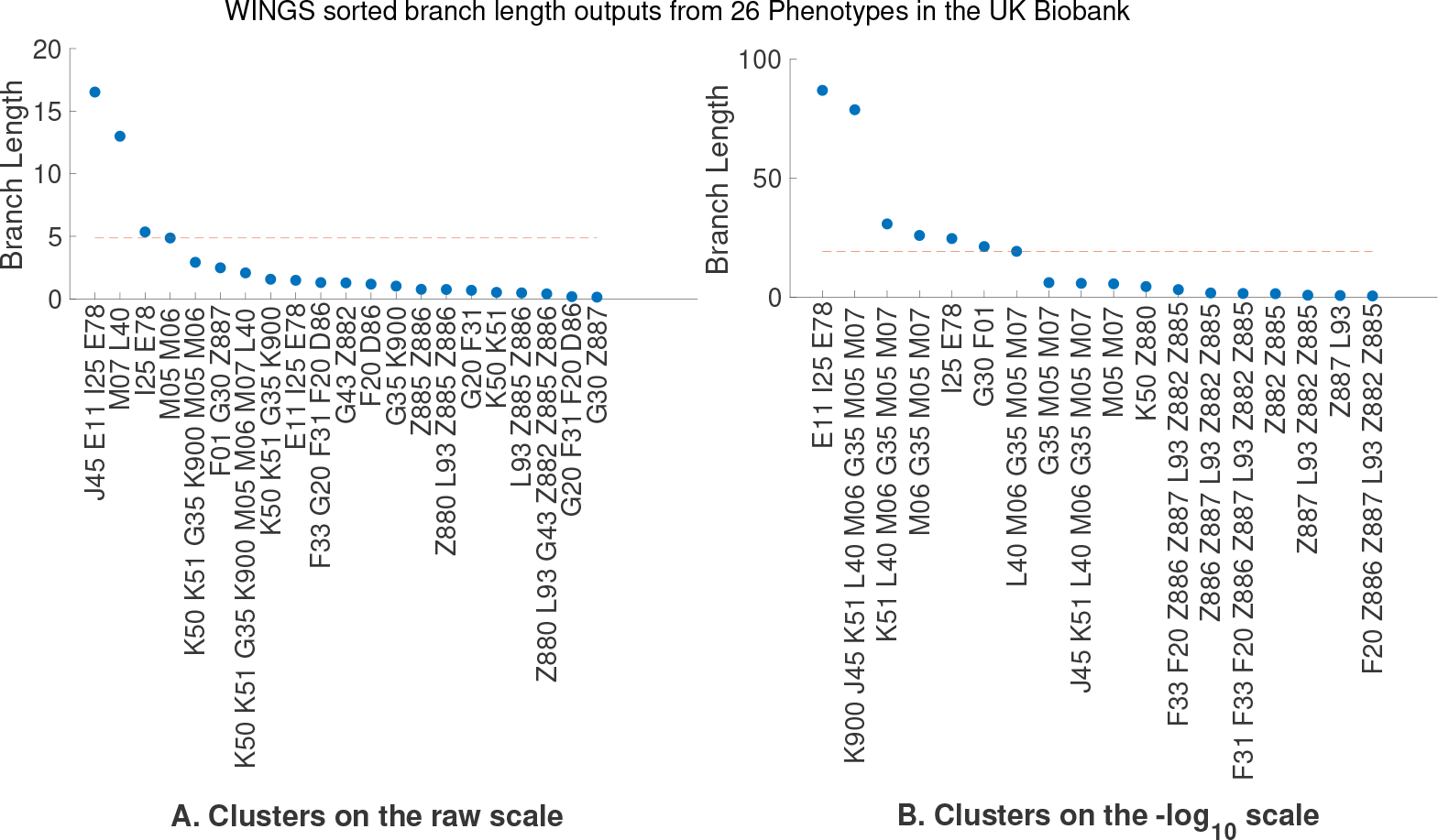
WINGS sorted branch lengths applied to 26 binary chronic illness phenotypes from the UK Biobank on the (A) raw and (B) −log_10_ scales. The sorted branch lengths corresponding to the branches in the dendrogram output of WINGS applied to the raw PEGASUS gene-level *p*-values (A) and −log_10_-transformed PEGASUS gene scores (B) for 26 case-control phenotypes in the UK Biobank. The dashed red horizontal line corresponds to the branch length threshold, where the identified significant clusters are those lying above the dashed line (boxed). Here, the x-axis shows the ICD10 codes; see Table S1 for the corresponding common disease names.

### 5.1 WINGS sensitivity to other gene level-association statistics

To showcase how WINGS can be used with any gene level association statistic we designed a similar set of simulations as outlined in 2.4 for two additional methods PASCAL (sum) [23], shown in Table S2, and SKAT [43], shown in Table S3.

**Figure S7:**
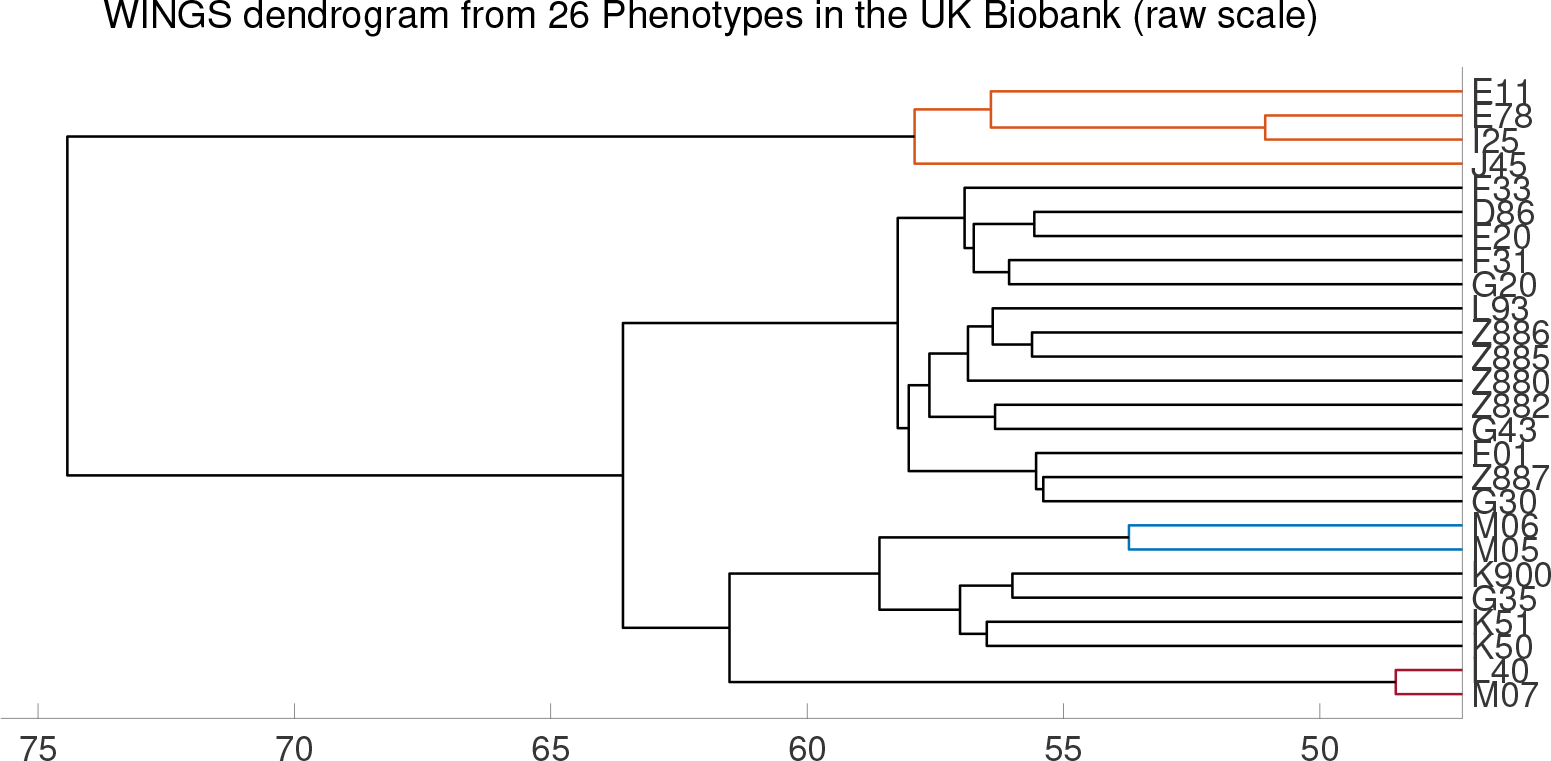
WINGS dendrogram applied to raw PEGASUS scores for 26 binary chronic illness phenotypes from the UK Biobank. The dendrogram output of WINGS to the raw PEGASUS scores of the 26 binary chronic illness phenotypes from the UK Biobank data. The color coded branches correspond to significant clusters identified by WINGS. The corresponding sorted branch lengths are presented in Figure S6(A) in the paper.

**Figure S8:**
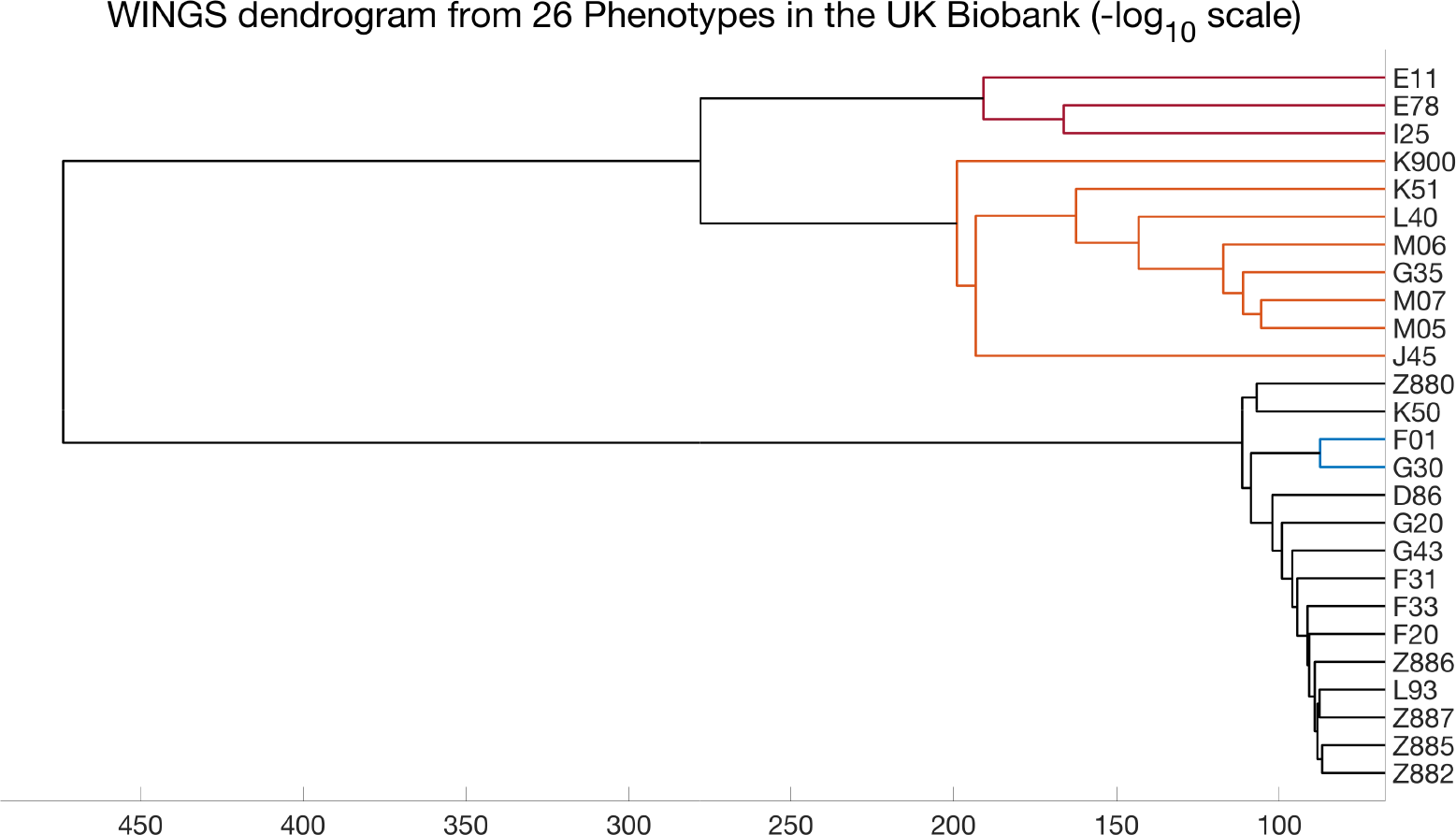
WINGS dendrogram applied to −log_10_ transformed PEGASUS scores for 26 binary chronic illness phenotypes from the UK Biobank. The dendrogram output of WINGS to the −log_10_ transformed PEGASUS scores of the 26 binary chronic illness phenotypes from the UK Biobank data. The color coded branches correspond to significant clusters identified by WINGS. The corresponding sorted branch lengths are presented in Figure S6(B) in the paper

**Figure S9:**
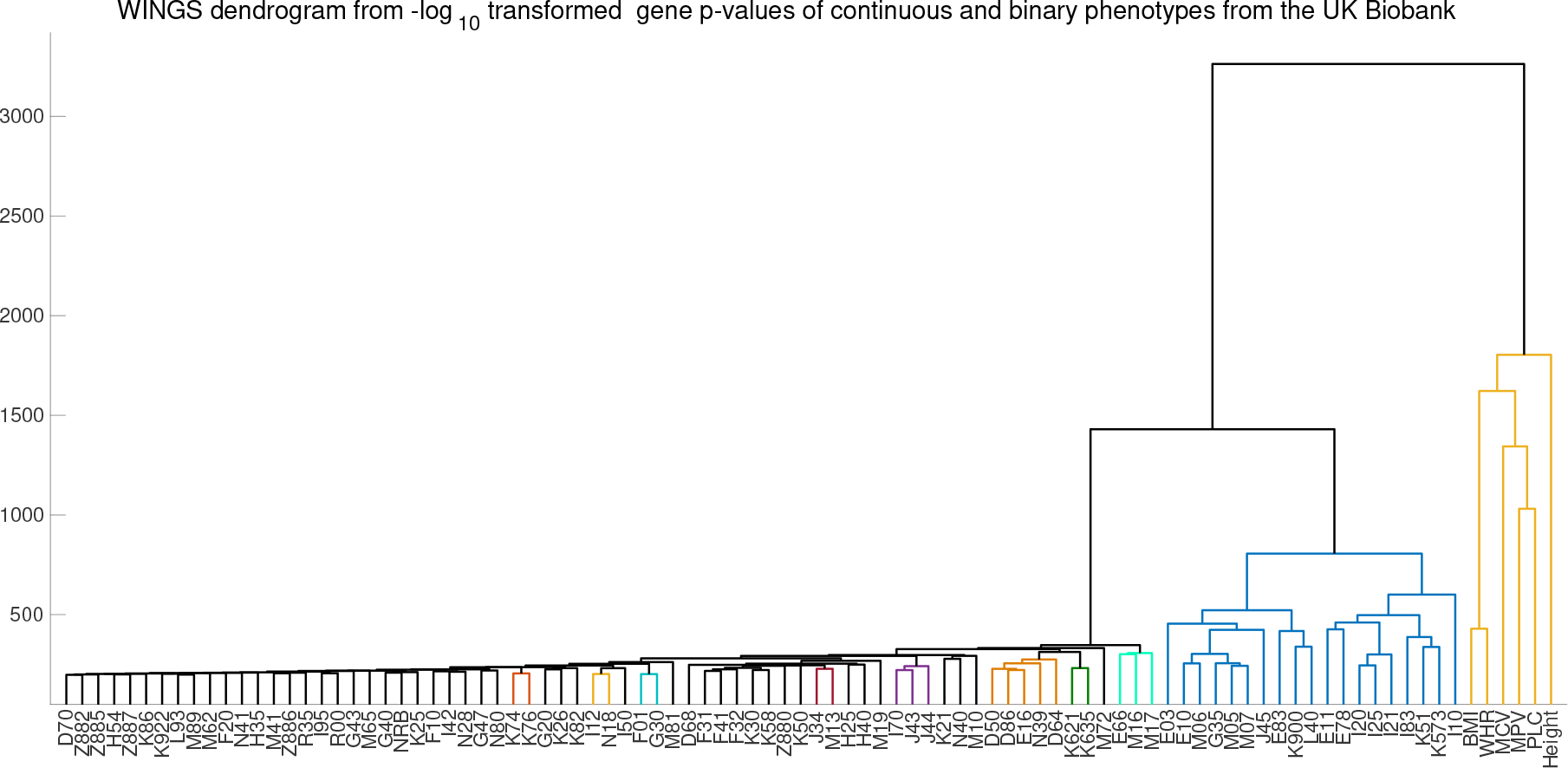
WINGS dendrogram from 94 case-control phenotypes in the UK Biobank separates continuous and binary traits. The dendrogram output of Ward hierarchical clustering applied to the −log_10_ transformed PEGASUS scores of the empirical continuous and binary traits. The branches are color coded by the largest significant clusters identified by the branch thresholding algorithm. The continuous phenotypes cluster together on the right of the dendrogram (in yellow), remaining disjoint from the remaining binary phenotypes until there is a single cluster.

**Figure S10:**
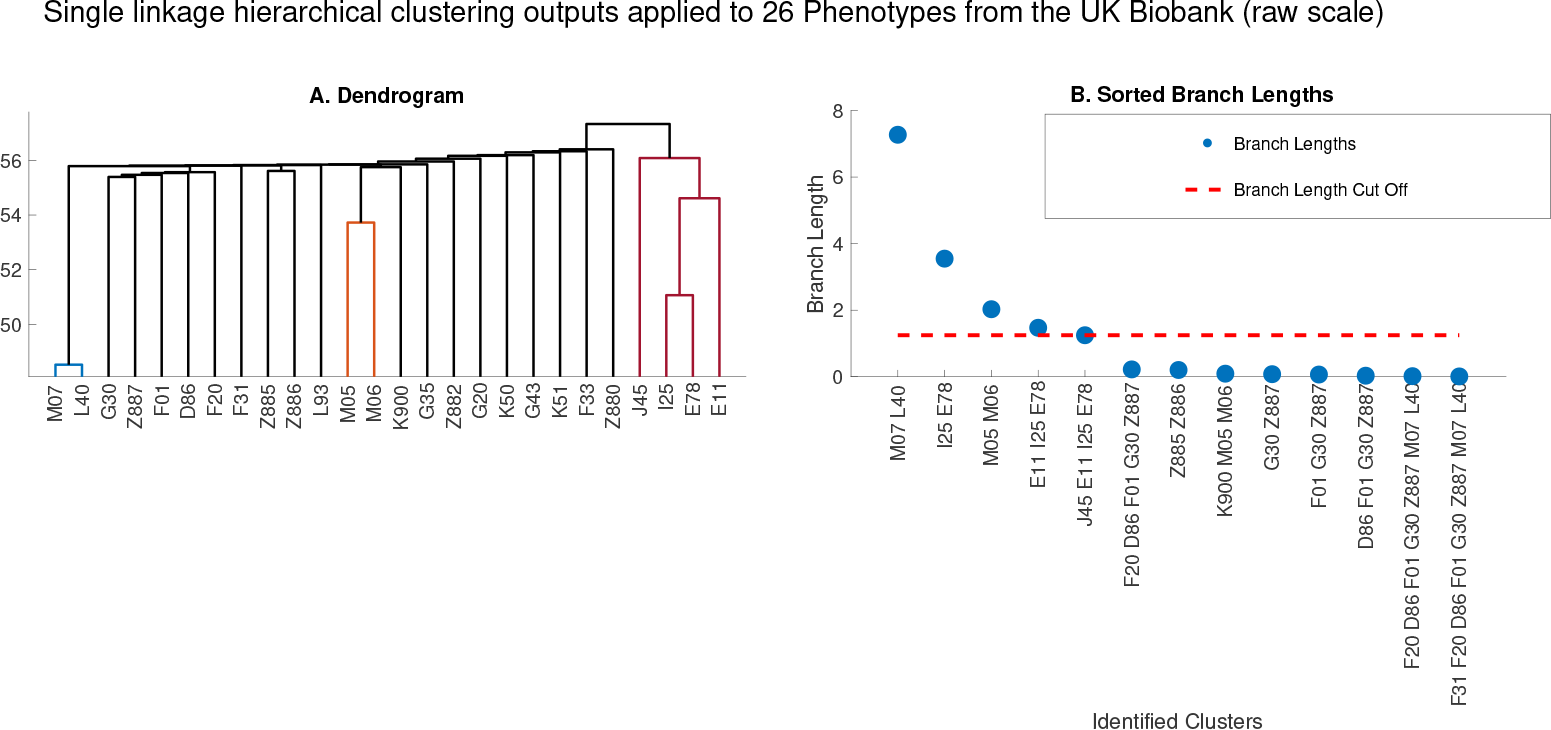
Single linkage clustering applied to PEGASUS *p*-values of 26 phenotypes from the UK Biobank (raw scale). (A) The dendrogram and (B) sorted branch lengths corresponding to the output of single linkage hierarchical clustering applied to the raw PEGASUS scores of the 26 phenotypes from the UK Biobank. The dashed red horizontal line on the right figure corresponds to the branch length threshold, where the identified significant clusters are those lying above the dashed line.

**Figure S11:**
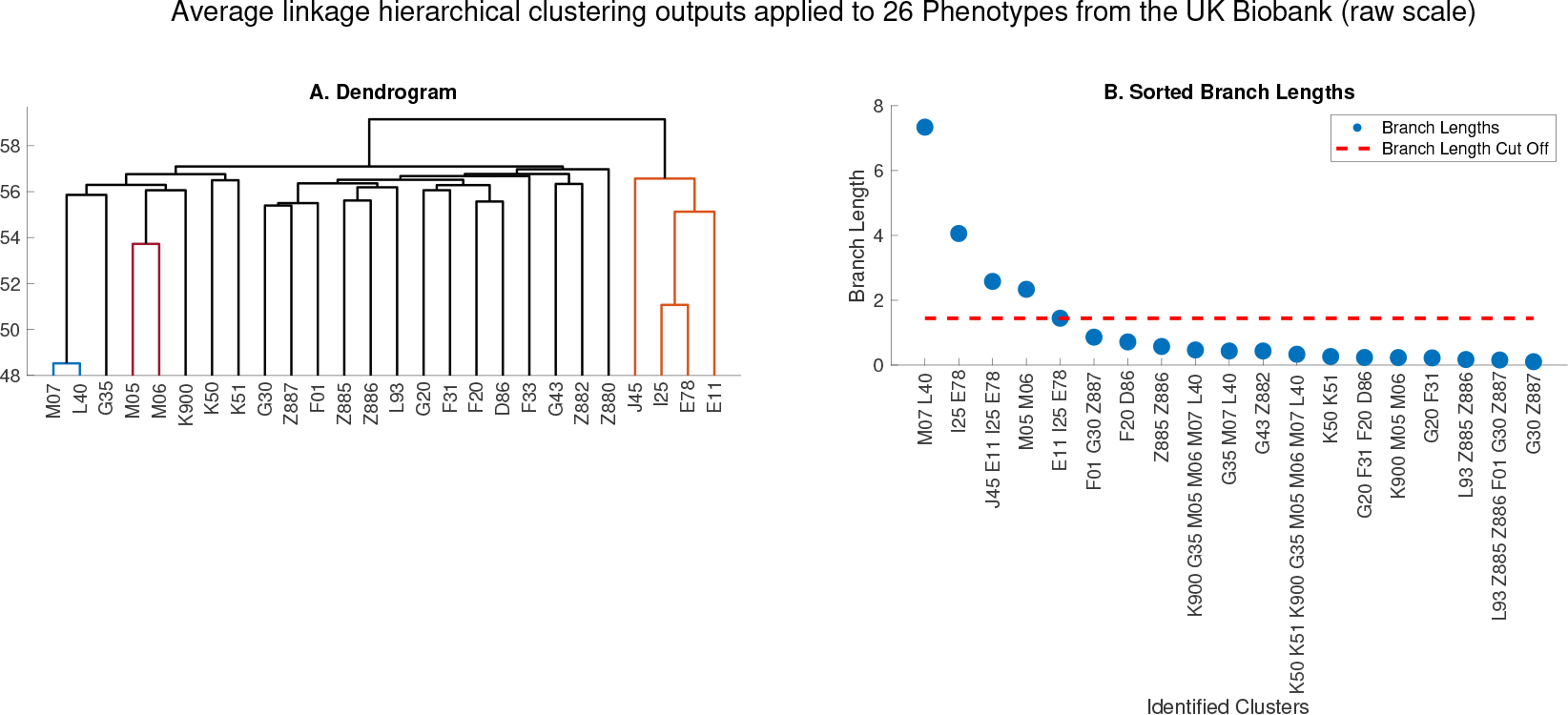
Average linkage clustering applied to PEGASUS *p*-values of 26 phenotypes from the UK Biobank (raw scale). (A) The dendrogram and (B) sorted branch lengths corresponding to the output of average linkage hierarchical clustering applied to the raw PEGASUS scores of the 26 phenotypes from the UK Biobank. The dashed red horizontal line on the right figure corresponds to the branch length threshold, where the identified significant clusters are those lying above the dashed line.

**Figure S12:**
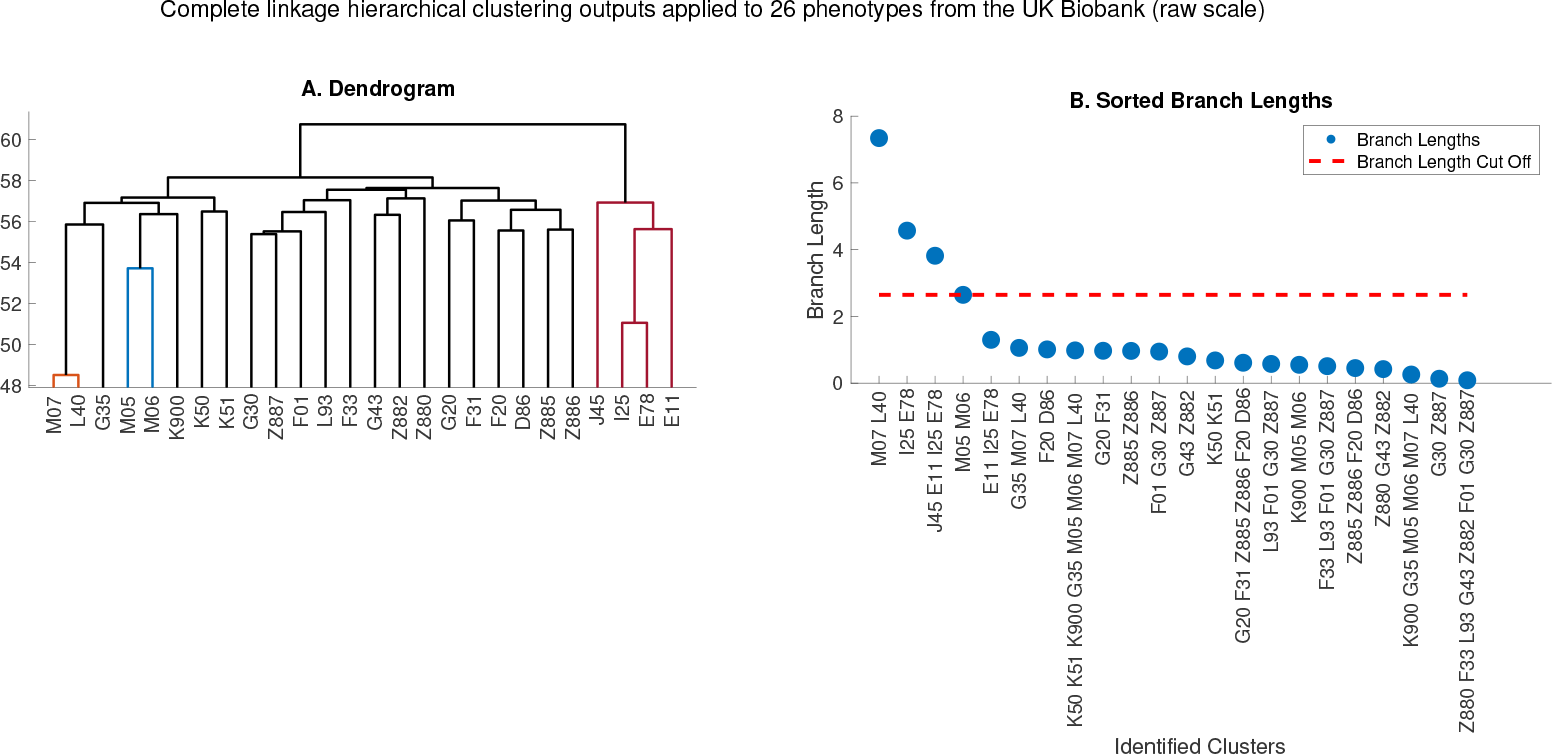
Complete linkage clustering applied to PEGASUS *p*-values of 26 phenotypes from the UK Biobank (raw scale). (A) The dendrogram and (B) sorted branch lengths corresponding to the output of complete linkage hierarchical clustering applied to the raw PEGASUS scores of the 26 phenotypes from the UK Biobank. The dashed red horizontal line on the right figure corresponds to the branch length threshold, where the identified significant clusters are those lying above the dashed line.

**Figure S13:**
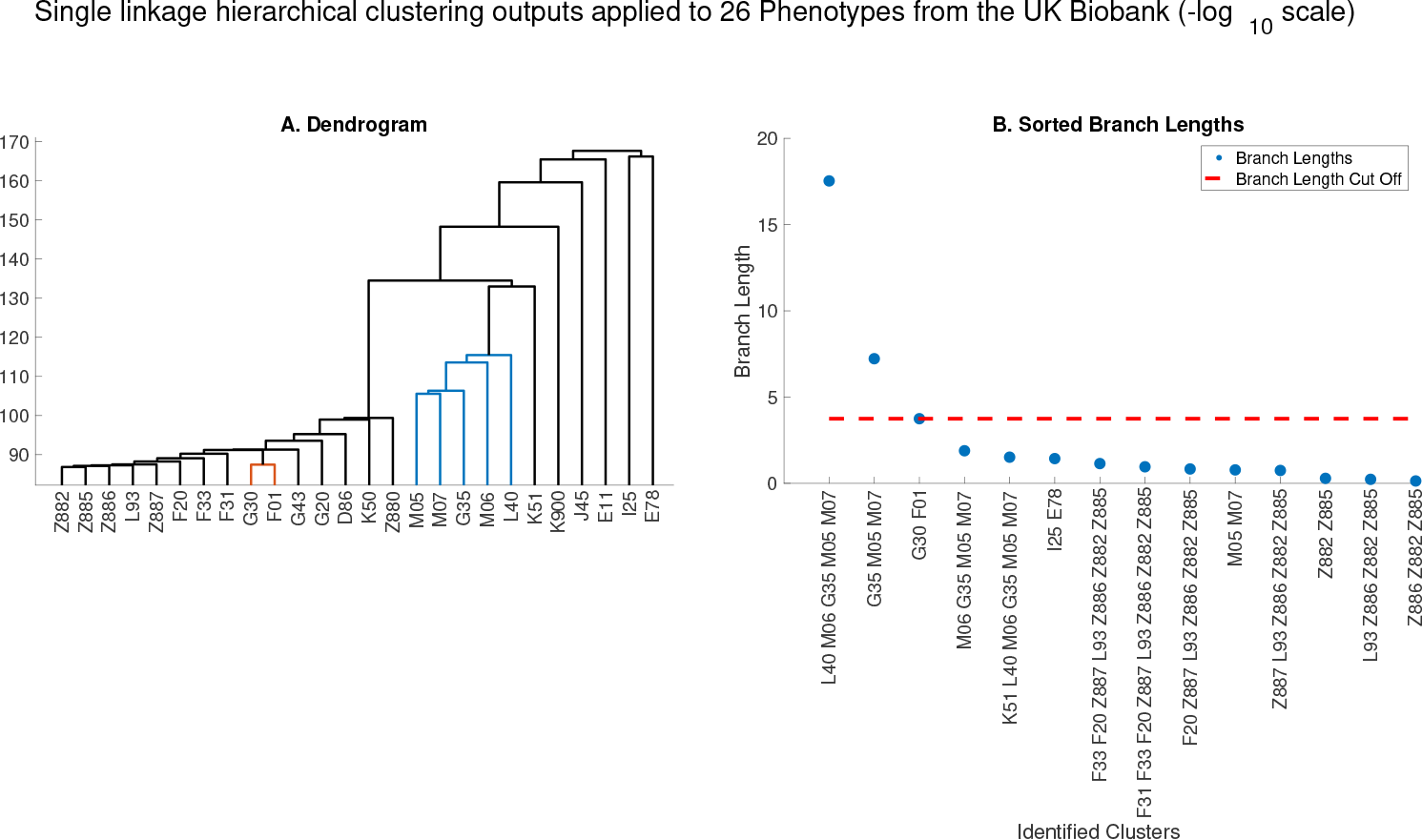
Single linkage clustering applied to −log_10_ transformed PEGASUS *p*-values of 26 phenotypes from the UK Biobank. (A) The dendrogram and (B) sorted branch lengths corresponding to the output of single linkage hierarchical clustering applied to the −log_10_ transformed PEGASUS scores of the 26 phenotypes from the UK Biobank. The dashed red horizontal line on the right figure corresponds to the branch length threshold, where the identified significant clusters are those lying above the dashed line.

**Figure S14:**
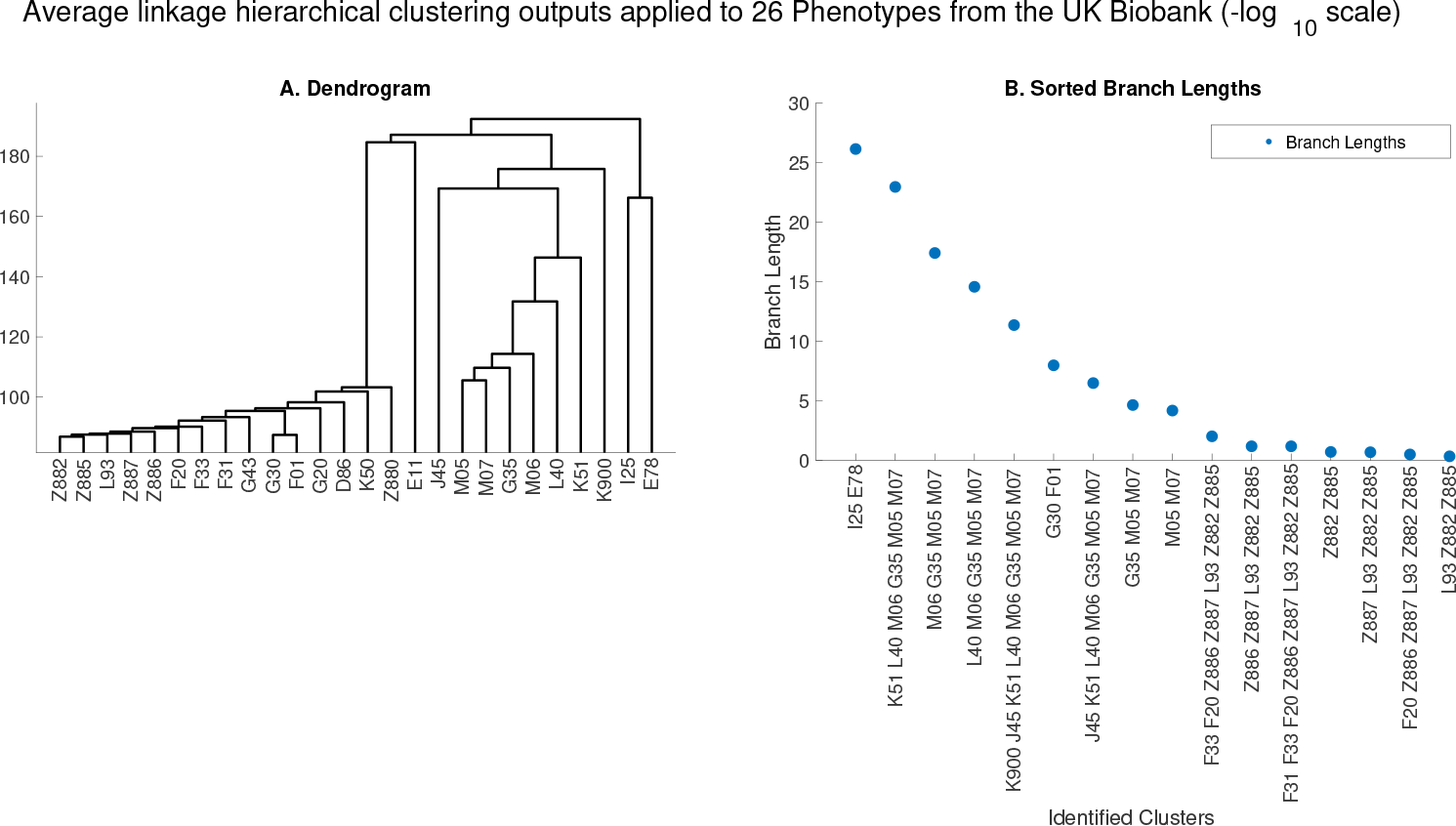
Average linkage clustering applied to −log_10_ transformed PEGASUS *p*-values of 26 phenotypes from the UK Biobank. (A) The dendrogram and (B) sorted branch lengths corresponding to the output of average linkage hierarchical clustering applied to the −log_10_ transformed PEGASUS scores of the 26 phenotypes from the UK Biobank. Here, there is no significant branch length threshold and consequently there are no significant clusters.

**Figure S15:**
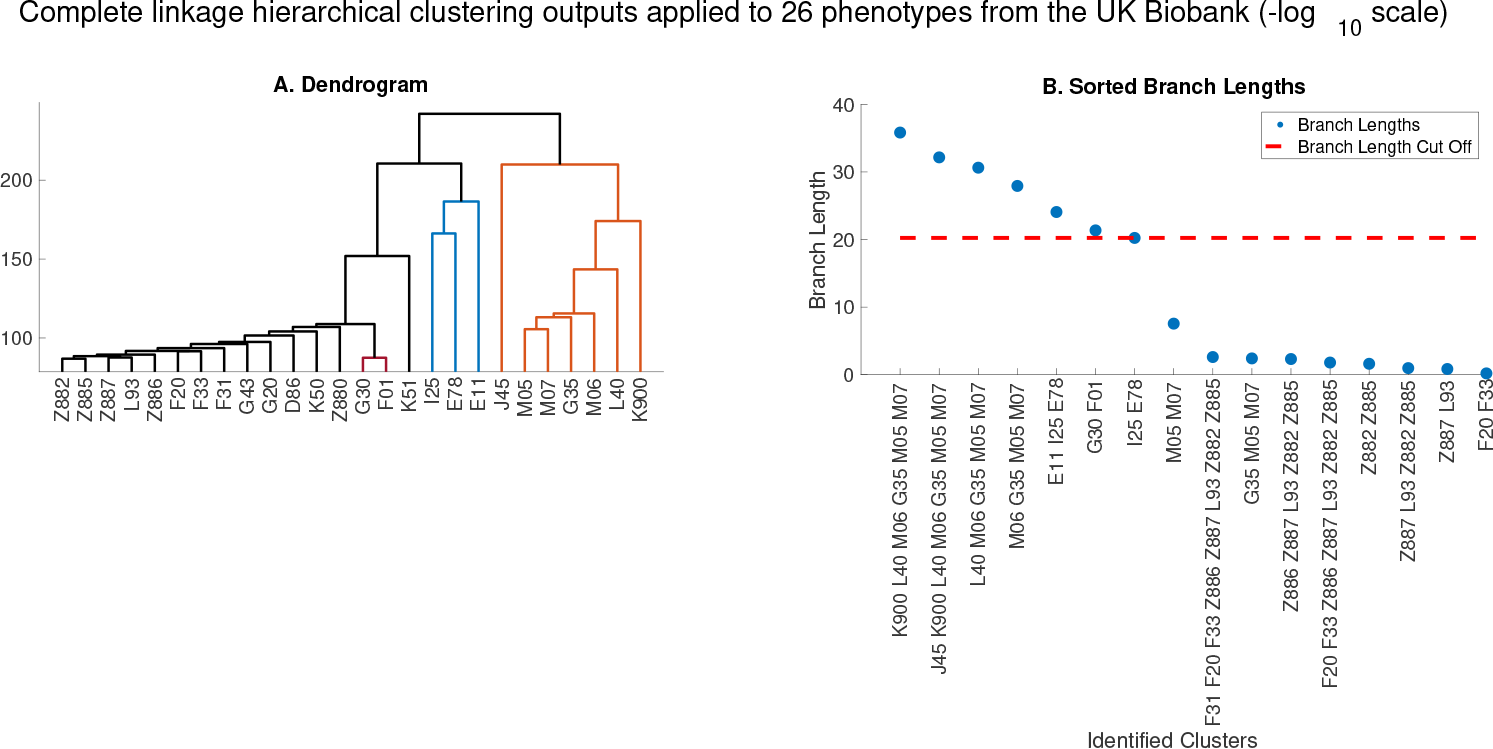
Complete linkage clustering applied to −log_10_ transformed PEGASUS *p*-values of 26 phenotypes from the UK Biobank. (A) The dendrogram and (B) sorted branch lengths corresponding to the output of complete linkage hierarchical clustering applied to the −log_10_ transformed PEGASUS scores of the 26 phenotypes from the UK Biobank. The dashed red horizontal line on the right figure corresponds to the branch length threshold, where the identified significant clusters are those lying above the dashed line.

**Figure S16:**
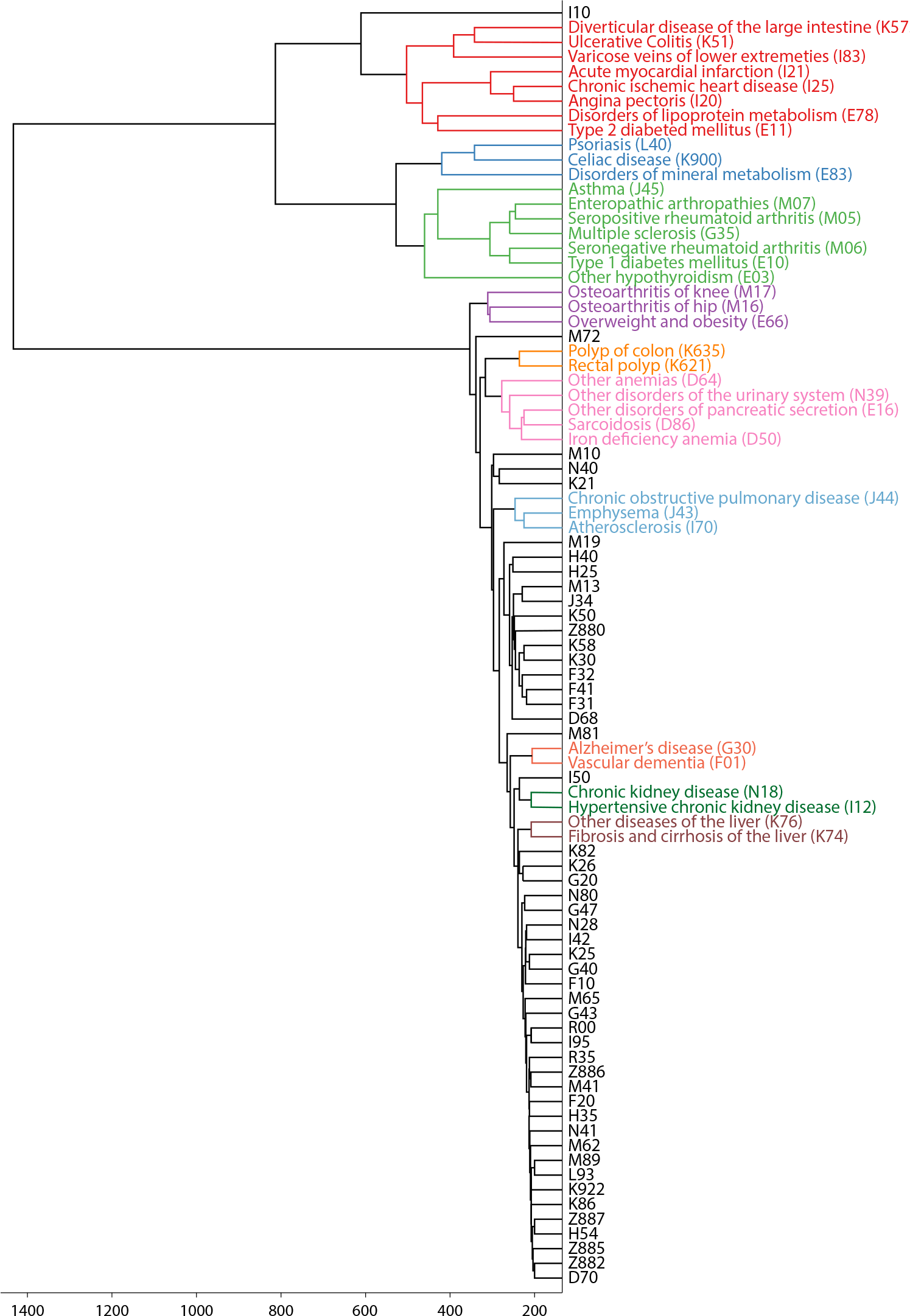
WINGS dendrogram from 87 case-control phenotypes using both genes and intergenic regions as features. We analyzed a matrix of PEGASUS *p*-values on the −log_10_ scale using both genes and intergenic regions as features. The topology of the tree is highly preserved compared to the dendrogram shown in Figure 4.

## References

[1] Farhad Hormozdiari, Gleb Kichaev, Wen-Yun Yang, Bogdan Pasaniuc, and Eleazar Eskin. Identification of causal genes for complex traits. Bioinformatics, 31(12):206–213, 2015.

[2] Huwenbo Shi, Gleb Kichaev, and Bogdan Pasaniuc. Contrasting the genetic architecture of 30 complex traits from summary association data. The American Journal of Human Genetics, 99(1):139–153, 2016.

[3] Evan A Boyle, Yang I Li, and Jonathan K Pritchard. An expanded view of complex traits: from polygenic to omnigenic. Cell, 169(7):1177–1186, 2017.

[4] Marcus W Feldman and Sohini Ramachandran. Missing compared to what? revisiting heritability, genes and culture. Phil. Trans. R. Soc. B, 373(1743):20170064, 2018.

[5] JE Huffman. Examining the current standards for genetic discovery and replication in the era of mega-biobanks. Nature communications, 9(1):5054, 2018.

[6] Evangelos Evangelou, Helen R Warren, David Mosen-Ansorena, Borbala Mifsud, Raha Pazoki, He Gao, Georgios Ntritsos, Niki Dimou, Claudia P Cabrera, Ibrahim Karaman, et al. Genetic analysis of over 1 million people identifies 535 new loci associated with blood pressure traits. Nature genetics, 50(10):1412, 2018.

[7] Yan Zhang, Guanghao Qi, Ju-Hyun Park, and Nilanjan Chatterjee. Estimation of complex effectsize distributions using summary-level statistics from genome-wide association studies across 32 complex traits. Nature genetics, 50(9):1318, 2018.

[8] Clare Bycroft, Colin Freeman, Desislava Petkova, Gavin Band, Lloyd T Elliott, Kevin Sharp, Allan Motyer, Damjan Vukcevic, Olivier Delaneau, Jared O’Connell, et al. Genome-wide genetic data on∼ 500,000 uk biobank participants. BioRxiv, page 166298, 2017.

[9] Dan M Roden, Jill M Pulley, Melissa A Basford, Gordon R Bernard, Ellen W Clayton, Jeffrey R Balser, and Dan R Masys. Development of a large-scale de-identified dna biobank to enable personalized medicine. Clinical Pharmacology & Therapeutics, 84(3):362–369, 2008.

[10] Joshua C Denny, Marylyn D Ritchie, Melissa A Basford, Jill M Pulley, Lisa Bastarache, Kristin Brown-Gentry, Deede Wang, Dan R Masys, Dan M Roden, and Dana C Crawford. Phewas: demonstrating the feasibility of a phenome-wide scan to discover gene–disease associations. Bioinformatics, 26(9):1205–1210, 2010.

[11] Joseph K Pickrell, Tomaz Berisa, Jimmy Z Liu, Laure Ségurel, Joyce Y Tung, and David A Hinds. Detection and interpretation of shared genetic influences on 42 human traits. Nature genetics, 48(7):709, 2016.

[12] Farhad Hormozdiari, Martijn van de Bunt, Ayellet V Segrè, Xiao Li, Jong Wha J Joo, Michael Bilow, Jae Hoon Sul, Sriram Sankararaman, Bogdan Pasaniuc, and Eleazar Eskin. Colocalization of gwas and eqtl signals detects target genes. The American Journal of Human Genetics, 99(6):1245–1260, 2016.

[13] Joshua C Denny, Lisa Bastarache, Marylyn D Ritchie, Robert J Carroll, Raquel Zink, Jonathan D Mosley, Julie R Field, Jill M Pulley, Andrea H Ramirez, Erica Bowton, et al. Systematic comparison of phenome-wide association study of electronic medical record data and genome-wide association study data. Nature biotechnology, 31(12):1102, 2013.

[14] Joshua C Denny, Lisa Bastarache, and Dan M Roden. Phenome-wide association studies as a tool to advance precision medicine. Annual review of genomics and human genetics, 17:353–373, 2016.

[15] Changjian Jiang and Zhao-Bang Zeng. Multiple trait analysis of genetic mapping for quantitative trait loci. Genetics, 140(3):1111–1127, 1995.

[16] Jonathan Marchini, Bryan Howie, Simon Myers, Gil McVean, and Peter Donnelly. A new multipoint method for genome-wide association studies by imputation of genotypes. Nature genetics, 39(7):906, 2007.

[17] Manuel AR Ferreira and Shaun M Purcell. A multivariate test of association. Bioinformatics, 25(1):132–133, 2008.

[18] Matthew Stephens. A unified framework for association analysis with multiple related phenotypes. PloS one, 8(7):e65245, 2013.

[19] Patrick Turley, Raymond K Walters, Omeed Maghzian, Aysu Okbay, James J Lee, Mark Alan Fontana, Tuan Anh Nguyen-Viet, Robbee Wedow, Meghan Zacher, Nicholas A Furlotte, et al. Multi-trait analysis of genome-wide association summary statistics using mtag. Nature genetics, 50(2):229, 2018.

[20] Peter M Visscher, Sarah E Medland, Manuel AR Ferreira, Katherine I Morley, Gu Zhu, Belinda K Cornes, Grant W Montgomery, and Nicholas G Martin. Assumption-free estimation of heritability from genome-wide identity-by-descent sharing between full siblings. PLoS genetics, 2(3):e41, 2006.

[21] Jimmy Z Liu, Allan F Mcrae, Dale R Nyholt, Sarah E Medland, Naomi R Wray, Kevin M Brown, Nicholas K Hayward, Grant W Montgomery, Peter M Visscher, Nicholas G Martin, et al. A versatile gene-based test for genome-wide association studies. The American Journal of Human Genetics, 87(1):139–145, 2010.

[22] Michael C Wu, Seunggeun Lee, Tianxi Cai, Yun Li, Michael Boehnke, and Xihong Lin. Rare-variant association testing for sequencing data with the sequence kernel association test. The American Journal of Human Genetics, 89(1):82–93, 2011.

[23] David Lamparter, Daniel Marbach, Rico Rueedi, Zoltán Kutalik, and Sven Bergmann. Fast and rigorous computation of gene and pathway scores from snp-based summary statistics. PLoS computational biology, 12(1):e1004714, 2016.

[24] Or Zuk, Eliana Hechter, Shamil R Sunyaev, and Eric S Lander. The mystery of missing heritability: Genetic interactions create phantom heritability. Proceedings of the National Academy of Sciences, 109(4):1193–1198, 2012.

[25] Diana Chang and Alon Keinan. Principal component analysis characterizes shared pathogenetics from genome-wide association studies. PLoS computational biology, 10(9):e1003820, 2014.

[26] Priyanka Nakka, Benjamin J Raphael, and Sohini Ramachandran. Gene and network analysis of common variants reveals novel associations in multiple complex diseases. Genetics, 204(2):783–798, 2016.

[27] Priyanka Nakka, Natalie P Archer, Heng Xu, Philip J Lupo, Benjamin J Raphael, Jun J Yang, and Sohini Ramachandran. Novel gene and network associations found for lymphoblastic leukemia using case-control and family-based studies in multi-ethnic populations. Cancer Epidemiology and Prevention Biomarkers, pages cebp–0360, 2017.

[28] Nicola Aceto, Aditya Bardia, David T. Miyamoto, Maria C. Donaldson, Ben S. Wittner, Joel A. Spencer, Min Yu, Adam Pely, Amanda Engstrom, Huili Zhu, Brian W. Brannigan, Ravi Kapur, Shannon L. Stott, Toshi Shioda, Sridhar Ramaswamy, David T. Ting, Charles P. Lin, Mehmet Toner, Daniel A. Haber, and Shyamala Maheswaran. Circulating tumor cell clusters are oligoclonal precursors of breast cancer metastasis. Cell, 158(5):1110–1122, 2014.

[29] Jessica C.S. Brown, Justin Nelson, Benjamin VanderSluis, Raamesh Deshpande, Arielle Butts, Sarah Kagan, Itzhack Polacheck, Damian J. Krysan, Chad L. Myers, and Hiten D. Madhani. Unraveling the biology of a fungal meningitis pathogen using chemical genetics. Cell, 159(5):1168–1187, 2014.

[30] Inti A. Pagnuco, Juan I. Pastore, Guillermo Abras, Marcel Brun, and Virginia L. Ballarin. Analysis of genetic association using hierarchical clustering and cluster validation indices. Genomics, 109(5):438–445, 2017.

[31] Peter Langfelder, Bin Zhang, and Steve Horvath. Defining clusters from a hierarchical cluster tree: the dynamic tree cut package for r. Bioinformatics, 24(5):719–720, 2008.

[32] Antoine E. Zambelli. A data-driven approach to estimating the number of clusters in hierarchical clustering. ISCB Comm J, 5(2809), 2016.

[33] Trevor Hastie, Robert Tibshirani, and Jerome Friedman. The Elements of Statistical Learning. Springer, 2009.

[34] Christopher C Chang, Carson C Chow, Laurent CAM Tellier, Shashaank Vattikuti, Shaun M Purcell, and James J Lee. Second-generation plink: rising to the challenge of larger and richer datasets. Gigascience, 4(1):7, 2015.

[35] Ani Manichaikul, Josyf C Mychaleckyj, Stephen S Rich, Kathy Daly, Michèle Sale, and WeiMin Chen. Robust relationship inference in genome-wide association studies. Bioinformatics, 26(22):2867–2873, 2010.

[36] Gad Abraham, Yixuan Qiu, and Michael Inouye. Flashpca2: principal component analysis of biobank-scale genotype datasets. Bioinformatics, 33(17):2776–2778, 2017.

[37] Joe H. Ward Jr. Hierarchical grouping to optimize an objective function. Journal of the American Statistical Association, 58(301):236–244, 1963.

[38] Joe H. Ward, Jr. and Marion E. Hook. Application of an hierarchical grouping procedure to a problem of grouping profiles. Educational and Psychological Measurement, 23(1):69–81, 1963.

[39] Bipul Hossen, Siraj-Ud Doulah, and Aminul Hoque. Methods for evaluating agglomerative hierarchical clustering for gene expression data: A comparative study. Computational Biology and Bioinformatics, 3(6):88–94, 2015.

[40] Laura Ferreira and David B. Hitchcock. A comparison of hierarchical methods for clustering functional data. Communications in Statistics Simulation and Computation, 38(9):1925–1949, 2009.

[41] Genevieve L Wojcik, WH Linda Kao, and Priya Duggal. Relative performance of gene-and pathwaylevel methods as secondary analyses for genome-wide association studies. BMC genetics, 16(1):34, 2015.

[42] Maxim V Kuleshov, Matthew R Jones, Andrew D Rouillard, Nicolas F Fernandez, Qiaonan Duan, Zichen Wang, Simon Koplev, Sherry L Jenkins, Kathleen M Jagodnik, Alexander Lachmann, et al. Enrichr: a comprehensive gene set enrichment analysis web server 2016 update. Nucleic acids research, 44(W1):W90–W97, 2016.

[43] Iuliana Ionita-Laza, Seunggeun Lee, Vlad Makarov, Joseph D Buxbaum, and Xihong Lin. Sequence kernel association tests for the combined effect of rare and common variants. The American Journal of Human Genetics, 92(6):841–853, 2013.

[44] Isabella Morlini and Sergio Zani. Dissimilarity and similarity measures for comparing dendrograms and their applications. Advances in Data Analysis and Classification, 6(2):85–105, Jul 2012.

[45] Edward Y Chen, Christopher M Tan, Yan Kou, Qiaonan Duan, Zichen Wang, Gabriela Vaz Meirelles, Neil R Clark, and Avi Maayan. Enrichr: interactive and collaborative html5 gene list enrichment analysis tool. BMC bioinformatics, 14(1):128, 2013.

[46] Maxim V Kuleshov, Matthew R Jones, Andrew D Rouillard, Nicolas F Fernandez, Qiaonan Duan, Zichen Wang, Simon Koplev, Sherry L Jenkins, Kathleen M Jagodnik, Alexander Lachmann, et al. Enrichr: a comprehensive gene set enrichment analysis web server 2016 update. Nucleic acids research, 44(W1):W90–W97, 2016.

[47] Sasha Bozeat, Carol A Gregory, Matthew A Lambon Ralph, and John R Hodges. Which neuropsychiatric and behavioural features distinguish frontal and temporal variants of frontotemporal dementia from alzheimer’s disease? Journal of Neurology, Neurosurgery & Psychiatry, 69(2):178–186, 2000.

[48] Richard J Perrin, Anne M Fagan, and David M Holtzman. Multimodal techniques for diagnosis and prognosis of alzheimer’s disease. Nature, 461(7266):916, 2009.

[49] Matthew Stephens. False discovery rates: a new deal. Biostatistics, 18(2):275–294, 2016.

[50] George Hripcsak, Matthew E Levine, Ning Shang, and Patrick B Ryan. Effect of vocabulary mapping for conditions on phenotype cohorts. Journal of the American Medical Informatics Association, 25(12):1618–1625, 2018.

[51] Chaitanya Shivade, Preethi Raghavan, Eric Fosler-Lussier, Peter J Embi, Noemie Elhadad, Stephen B Johnson, and Albert M Lai. A review of approaches to identifying patient phenotype cohorts using electronic health records. Journal of the American Medical Informatics Association, 21(2):221–230, 2013.

[52] Alicia R Martin, Christopher R Gignoux, Raymond K Walters, Genevieve L Wojcik, Benjamin M Neale, Simon Gravel, Mark J Daly, Carlos D Bustamante, and Eimear E Kenny. Human demographic history impacts genetic risk prediction across diverse populations. The American Journal of Human Genetics, 100(4):635–649, 2017.

[53] Jian Yang, Beben Benyamin, Brian P McEvoy, Scott Gordon, Anjali K Henders, Dale R Nyholt, Pamela A Madden, Andrew C Heath, Nicholas G Martin, Grant W Montgomery, et al. Common snps explain a large proportion of the heritability for human height. Nature genetics, 42(7):565, 2010.

[54] Andrew R Wood, Tonu Esko, Jian Yang, Sailaja Vedantam, Tune H Pers, Stefan Gustafsson, Audrey Y Chu, Karol Estrada, Jian’an Luan, Zoltán Kutalik, et al. Defining the role of common variation in the genomic and biological architecture of adult human height. Nature genetics, 46(11):1173, 2014.

